# Quantifying the impact of gut microbiota on inflammation and hypertensive organ damage

**DOI:** 10.1101/2021.09.17.460671

**Authors:** Ellen G. Avery, Hendrik Bartolomaeus, Ariana Rauch, Chia-Yu Chen, Gabriele N’Diaye, Ulrike Löber, Theda U. P. Bartolomaeus, Raphaela Fritsche-Guenther, André F. Rodrigues, Dmitry Tsvetkov, Mihail Todiras, Joon-Keun Park, Lajos Markó, András Maifeld, Michael Bader, Stefan Kempa, Jennifer A. Kirwan, Sofia K. Forslund, Dominik N. Müller, Nicola Wilck

## Abstract

**Aims:** Hypertension (HTN) can lead to heart and kidney damage. The gut microbiota has been linked to HTN, although it is difficult to estimate its significance due to the variety of other features known to influence HTN. In the present study, we used germ-free (GF) and colonized (COL) littermate mice to quantify the impact of microbial colonization on organ damage in HTN.

**Methods and results:** Four-week-old male GF C57BL/6J littermates were randomized to remain GF or receive microbial colonization. HTN was induced by subcutaneous infusion with angiotensin (Ang) II (1.44mg/kg/d) and 1% NaCl in the drinking water; sham-treated mice served as control. Renal damage was exacerbated in GF mice, whereas cardiac damage was more comparable between COL and GF, suggesting that the kidney is more sensitive to microbial influence. Multivariate analysis revealed a larger effect of HTN in GF mice. Serum metabolomics demonstrated that the colonization status influences circulating metabolites relevant to HTN. Importantly, GF mice were deficient in anti-inflammatory fecal short-chain fatty acids (SCFA). Flow cytometry showed that the microbiome has an impact on the induction of anti-hypertensive myeloid-derived suppressor cells and pro-inflammatory Th17 cells in HTN. In vitro inducibility of Th17 cells was significantly higher for cells isolated from GF than conventionally raised mice.

**Conclusions:** Microbial colonization status of mice had potent effects on their phenotypic response to a hypertensive stimulus, and the kidney is a highly microbiota-susceptible target organ in HTN. The magnitude of the pathogenic response in GF mice underscores the role of the microbiome in mediating inflammation in HTN.

**Translation Perspective:** To assess the potential of microbiota-targeted interventions to prevent organ damage in hypertension, an accurate quantification of microbial influence is necessary. We provide evidence that the development of hypertensive organ damage is dependent on colonization status and suggest that a healthy microbiota provides anti-hypertensive immune and metabolic signals to the host. In the absence of normal symbiotic host-microbiome interactions, hypertensive damage to the kidney in particular is exacerbated. We suggest that hypertensive patients experiencing perturbations to the microbiota, which are common in CVD, may be at a greater risk for target-organ damage than those with a healthy microbiome.

## Introduction

Hypertension (HTN) is the leading risk factor for non-communicable diseases worldwide,^1^ and is known as a multifactorial disease, where complex mechanisms often co-occur to lead to a persistent increase in blood pressure (BP). Several studies have indicated that alterations in the composition and function of the intestinal microbiota may contribute to the burden of hypertensive disease.^2-6^ However, it is difficult to estimate the contribution of the microbiota, especially in human studies, where the added complexities of other contributing factors easily obstruct our understanding. The aim of our study is to understand the relative contribution that the microbiota has to the burden of hypertensive disease.

Mounting evidence suggests that inflammation is not only characteristic of hypertensive cardiovascular disease (CVD) but is causally linked to disease progression and severity.^3^ Components of both the innate and adaptive immune system have been implicated.^3^ T helper 17 (Th17) cells and Type 1 helper T cells (Th1) have been shown to be integrally interlinked with hypertensive disease, and have been demonstrated to exacerbate cardiac and renal damage.^5, 7, 8^ Moreover, myeloid derived suppressor cells (MDSC) derived from hypertensive mice were shown to have immunosuppressive properties, and upon adoptive transfer were able to mitigate BP increase in response to angiotensin II (Ang II) infusion.^9^ MDSC, Th17, and Th1 cells have each been shown in different settings to be influenced by the microbiota.^5, 9-11^

We and others could recently demonstrate the role of several anti-inflammatory microbial metabolites in HTN. Short-chain fatty acids (SCFA) such as acetate, propionate, and butyrate, are produced by gut microbiota through the fermentation of indigestible dietary fiber.^12^ Acetate has been shown to ameliorate hypertensive damage to the kidney and heart in mice.^13^ Our recent work elucidated the protective role of propionate in Ang II-induced inflammation and cardiovascular damage.^14^ Furthermore, low butyrate levels have been associated with worsened CVD in several models.^15^ In addition to SCFA, we have recently shown that a bacterially produced indole metabolite derived from tryptophan suppresses Th17-driven inflammation in salt-sensitive HTN.^5^ In contrast, metabolites of microbial origin can also exacerbate disease in some contexts. For example, pro-inflammatory metabolites like trimethylamine N-oxide (TMAO) and indoxyl sulfate (IS) have been shown to aggravate CVD.^16, 17^

To address our central aim, we utilized germ-free (GF) mice. C57BL/6J GF littermates were randomized at four-weeks of age to either receive a microbiota transfer from our in-house C57BL/6J colony, or to remain GF for the duration of the experiment. Here we have uncovered several differences between GF and colonized (COL) mice in response to Ang II and 1 % NaCl in the drinking water, which underscores the importance of the microbiota in the pathogenesis of HTN-induced organ damage. Of note, we show an exacerbation of damage in GF mice compared to COL mice, which is more distinct in the kidney than in the heart.

## Materials and Methods

*Detailed description of all analytical methods and data analysis used are available in the Supplemental Material*.

## Results

### Absence of microbiota exacerbates cardiorenal damage

To exclude confounding effects relating to the genetic background of mice used in our study, GF littermates were randomized at 4 weeks of age for colonization with SPF microbiota (COL) or further kept under GF conditions. At 12 weeks of age, we induced HTN by subcutaneous Ang II infusion and 1% NaCl-supplemented drinking water. After 14 days, we analyzed hypertensive target organ damage (Figure 1A). Of note, we did not include surgical uninephrectomy to avoid bacterial contamination. The standard model in our lab includes uninephrectomy to induce a more severe form of renal damage ^8, 18^, thus we expected a lower degree of renal damage when compared to the published literature.^19^ To validate the integrity of our experiment, we first checked the colonization status of the mice. For this, gross morphological changes were assessed, and the characteristics typical of the gastrointestinal (GI) tract in GF mice (e.g., megacecum) did not persist in the mice which had been colonized (COL) (Supplemental Figure S1A). To confirm the microbial status of the respective group, we examined fecal pellets produced on the final day of experimentation. First, pellets were incubated in a thioglycolate medium for 96 hours, and GF mice were found to show no bacterial growth (Supplemental Figure S1B). Second, 16S rDNA copies per gram stool measured by qPCR were found to be similar in COL (Sham and HTN) and conventional SPF mice (CONV), whereas GF mice did not have more 16S rDNA copies than blank samples (Supplemental Figure S1C). Lastly, serum metabolomics confirmed the presence of bacterially derived metabolites in COL mice only (Supplemental Figure S1D). Shotgun metagenomics revealed that the grafted bacteria in COL mice (Supplemental Figure S1E-F, Supplemental File S1-4) showed the largest overlap with mouse gut metagenomes published in the global microbial gene catalogue (GMGC) (Supplemental Figure S2). We therefore concluded that the GF and COL groups were maintained as intended to confidently proceed with further analyses.

**Figure 1.**
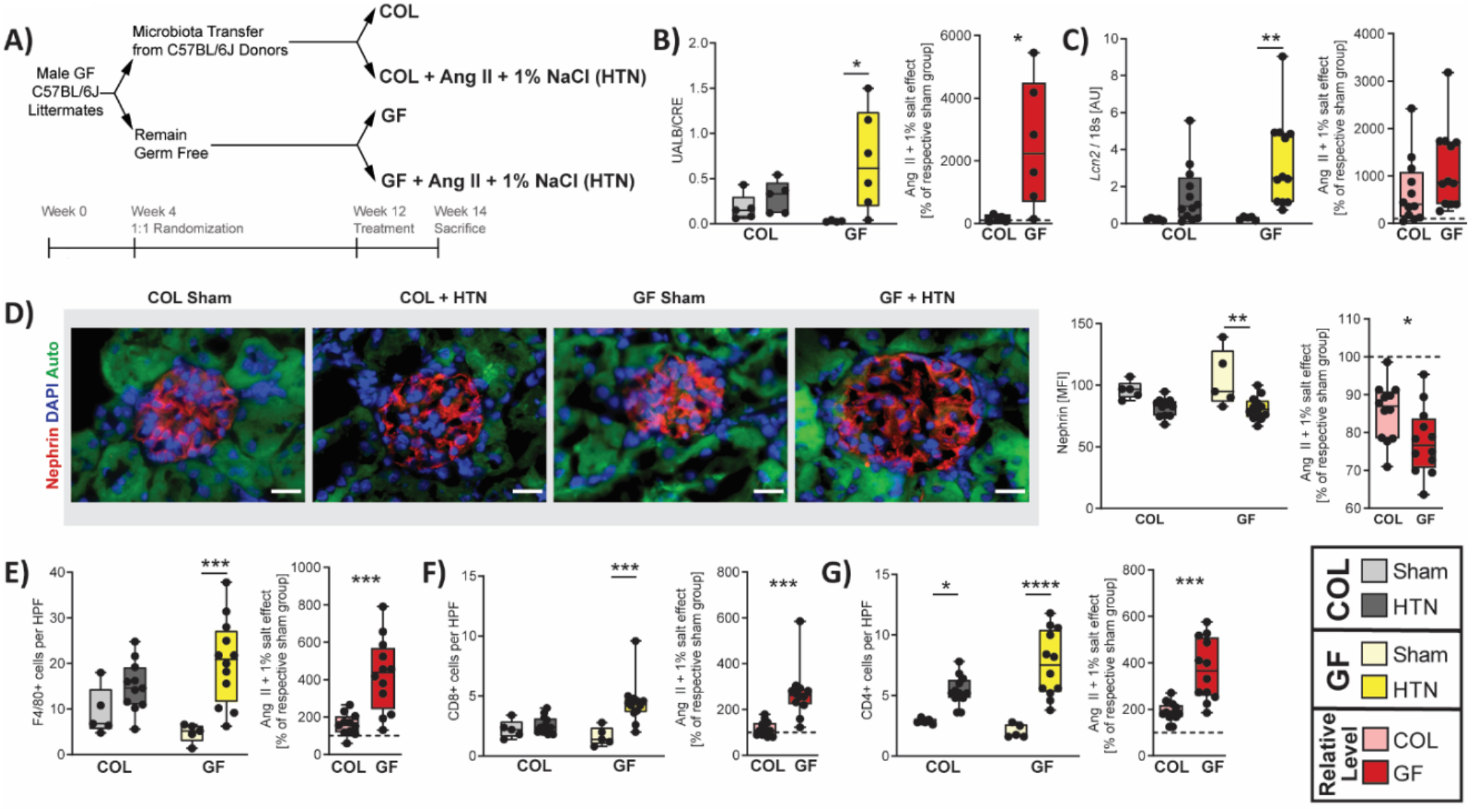
Renal damage is exacerbated under germ-free conditions. A) Description of experimental protocol (unless otherwise stated, GF Sham n = 5, GF + HTN n = 12, COL Sham n = 5, COL + HTN n = 12 for further analyses). B) Urinary albumin-to-creatinine ratio from spot urine collected upon sacrifice from a subset of mice (COL n = 5, COL + HTN n = 5, GF n = 4, GF + HTN n = 6). C) *Lcn2* gene expression was measured from kidney tissue by qPCR. D) Histological kidney sections were stained for nephrin. Representative glomeruli are shown (left). Nephrin immunofluorescence was quantified as mean fluorescence intensity (MFI) in the glomerular space averaged per mouse. The scale bar represents 20 µm. E) Macrophages (F4/80+), (F) CD8+, and (G) CD4+ T cells from kidney sections were counted from 5 representative high-power fields within the cortex. For (B-G), the left graph was tested using a two-way ANOVA and post-hoc Sidak multiple comparison’s test and depicts the raw values for each variable. For (B-F), hypertension was identified as the source of variation using two-way ANOVA, and post-hoc multiple comparison between sham and HTN within each group revealed that the GF comparison was the source of variation. In (G), hypertension was identified as the source of variation using two-way ANOVA, and post-hoc multiple comparison between sham and HTN within each group revealed that both the GF and COL comparison were significant. For B through G, the right plot depicts the relative change induced by Ang II + 1% NaCl in comparison to the respective sham group, tested using an unpaired two-tailed T-test. No change (100%) depicted as dotted line. For all plots, p-values are as follows; * P ≤ 0.05, ** P ≤ 0.01, *** P ≤ 0.001, **** P ≤ 0.0001.

One of the hallmarks of hypertensive target organ damage is renal damage, which is characterized by abnormally high excretion of albumin with the urine (albuminuria), fibrosis, and inflammation. In line with the literature^19^, our HTN induction without uninephrectomy lead to a moderate increase in albuminuria in COL mice (Figure 1B). GF mice developed a greater degree of albuminuria upon HTN induction, which is abundantly clear when comparing the relative increase of GF and COL mice compared to their respective sham groups (Figure 1B). HTN also lead to a significant increase in renal damage marker lipocalin-2 in GF mice (Lcn2), which was not evident in COL mice (Figure 1C). Next, we analyzed nephrin, a protein in the podocytes’ slit membrane, by immunofluorescence. We observed a significant decrease of nephrin immunofluorescence in GF mice, where COL mice exhibited a similar but insignificant trend (Figure 1D). HTN led to a significant increase of macrophages (F4/80+ cells, Figure 1E) and cytotoxic T cells (CD8+ cells, Figure 1F) in the kidney of GF mice, not reaching significance in COL mice. Likewise, we found that mRNA expression of CC-chemokine ligand 2 (Ccl2, Supplemental Figure S3A) and infiltrating T cells (CD3+, Supplemental Figure S3B) were selectively increased in the GF group upon HTN. While T helper cells (CD4+ cells) were shown to increase in both GF and COL mice, GF mice displayed a stronger increase (Figure 1G). The number of leukocytes (CD45+ cells) within the kidney confirms the stronger effect of HTN on renal inflammation in GF. (Supplemental Figure S3C). For all immune populations in the kidney, the change in HTN relative to sham was consistently exacerbated in GF compared to COL (Figure 1E-G, Supplemental Figure S3B-C). Lastly, we investigated kidney fibrosis. Expression of *Col3a1* was significantly increased only in GF+HTN mice (Supplemental Figure S3D). Perivascular fibrosis analyzed by Masson’s trichrome staining was accentuated in GF mice but not statistically different between the groups using two-way ANOVA; although when comparing the relative increase from sham to HTN, there was a significant difference between GF and COL (Supplemental Figure S3E). Similar to what was previously shown^20^, GF mice tended to have lower baseline values for several damage markers when comparing sham-treated GF and COL mice. Overall, renal pathology upon HTN induction was greater in GF mice when compared to their COL littermates.

Next, we examined the cardiac phenotype. To evaluate cardiac hypertrophy upon HTN induction we calculated the heart weight-to-tibia length ratio (Figure 2A), which indicated that there was more significant damage to the GF mice than COL in HTN. Left ventricular weight taken from echocardiography relative to the tibia length (Figure 2B) as well as cardiac *Nppb* expression (Figure 2C) confirmed this finding. Neither the GF+HTN nor COL+HTN mice had a reduced ejection fraction (Figure 2D), indicating none of these mice were experiencing systolic heart failure. Using two-way ANOVA, both perivascular (Figure 2E) and interstitial (Figure 2F) fibrosis were significantly increased in GF+HTN and not in COL+HTN mice compared to their respective sham group. Interestingly, when assessing the relative increase in HTN compared to sham for markers of cardiac fibrosis, there was no difference in GF compared to COL mice (Figure 2E-F). Next, we examined cardiac inflammation. Despite an increase in *Ccl2* expression in GF and not COL (Supplemental Figure S4A), macrophages (F4/80+) were increased in both GF and COL hearts upon HTN; and GF+HTN showed significantly less macrophages than COL+HTN mice (Figure 2G). The change in overall leukocytes (CD45+) within the heart mimics the changes seen for macrophages (Supplemental Figure S4B). Furthermore, no significance was reached when comparing CD4+ T helper cell infiltration (Figure 2H), whereas CD8+ cytotoxic T cells in the hearts increased in both GF+HTN and COL+HTN mice compared to sham (Figure 2I). We also observed an increase in the cardiac expression of pro-inflammatory cytokine *Tnfa* selectively in the GF+HTN group compared to sham (Supplemental Figure S4C). Altogether, cardiac hypertrophy and inflammation following HTN were affected to a greater extent in GF, but when assessing the relative change for GF and COL mice in HTN compared to sham we saw many similarities in the development of the cardiac fibrosis. Whereas in the kidney, there was a clear difference in the development of HTN damage between GF and COL mice, these distinctions were less evident in the heart.

**Figure 2.**
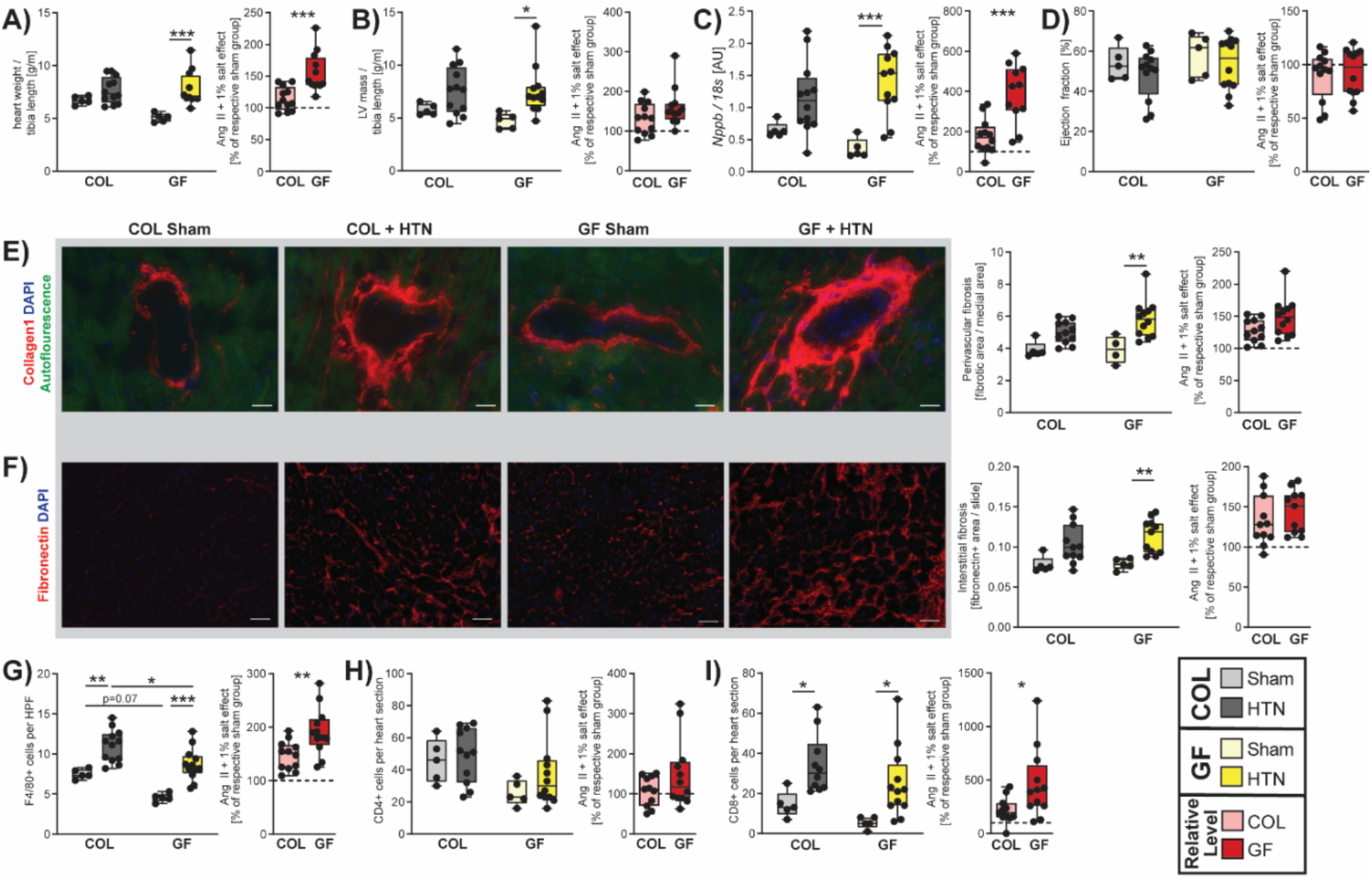
Cardiac inflammation and hypertrophy are aggravated in GF mice. A) Heart weight to tibia length [g/m] from mice taken at sacrifice. B) Left ventricular (LV) mass was estimated using echocardiography and normalized to tibia length [g/m]. C) Cardiac ventricular natriuretic peptide gene expression (*Nppb*) measured by qPCR. D) Echocardiography prior to sacrifice revealed no change in the ejection fraction upon HTN induction in GF or COL mice. E) Perivascular fibrosis from cardiac vessels, evaluated by measuring the fibrotic area relative to the medial area of vessels from Collagen 1 stained histology slides. F) Cardiac interstitial fibrosis was evaluated as fibronectin positive area proportionate to total area from 5 representative 40x magnification pictures from histological slides. In scale bar represents 40 µm. G) Macrophages (F4/80+) were counted from 5 representative high-power fields, and (H) CD4+, and (I) CD8+ T cells were counted from whole heart sections. Two-way ANOVA and post-hoc Sidak multiple comparison’s test for (A-I) was used to test significance in the left plot. In (A-C) and (E-F), hypertension was identified as the source of variation using two-way ANOVA, and post-hoc multiple comparison between sham and HTN within each group revealed that the GF comparison was the source of variation. In (G) hypertension and the microbiome were both identified as sources of variation using two-way ANOVA, and significant post-hoc comparisons are shown. In (I), hypertension was identified as the source of variation using two-way ANOVA, and post-hoc multiple comparison between sham and HTN within each group revealed that both the GF and COL comparison were significant. To the right in (A-I), the relative change induced by Ang II + 1% NaCl in comparison to the respective sham group was tested using an unpaired two-tailed T-test. No change (100%) depicted as dotted line. For all plots, p-values are as follows; * P ≤ 0.05, ** P ≤ 0.01, *** P ≤ 0.001.

### Hypertensive kidney damage is more sensitive to microbial status than cardiac damage

The aforementioned findings indicate that GF mice respond more sensitively to HTN. Within the context of our initial statistical approach (two-way ANOVA) we often saw a loss of significance for the HTN effect in COL mice under equal statistical power; and the differences between GF and COL response where much clearer when assessing the relative increase for a given marker in HTN (unpaired T-test). To expand on this idea to increase our understanding of the differences between GF and COL, we assessed the size of the HTN-induced effect by calculating an effect size (Cliff’s delta) and fold change for each marker. Using a comprehensive univariate testing strategy, we assessed the significance in the different tissue spaces using a robust false discovery rate (FDR) correction within GF and COL groups to root out any spurious findings. From the majority of kidney parameters assessed, across the subcategorizations of damage, fibrosis, and inflammatory markers, a very consistent pattern emerged in that the GF mice experienced worsened kidney outcomes compared to COL mice (Figure 3A, Supplemental Table 1). In contrast to the renal damage, both GF and COL mice experienced a more similar cardiac damage pattern, particularly regarding markers of cardiac fibrosis (Figure 3B, Supplemental Table 2). Albeit several cardiac parameters reached significance in GF and COL mice, the fold changes observed in GF mice were often larger (e.g., *Nppb*, perivascular fibrosis, *Ccn2, Lcn2*, F4/80, CD8; Supplemental Table 2). Interestingly, GF+HTN mice develop a significant increase in lung weight-to-tibia length ratio (Figure 3B), indicating the development of lung congestion^21^ due to aggravated cardiac dysfunction. Taken together, there was more overlap in the cardiac response to HTN in GF and COL groups than was seen for the kidney parameters.

**Figure 3.**
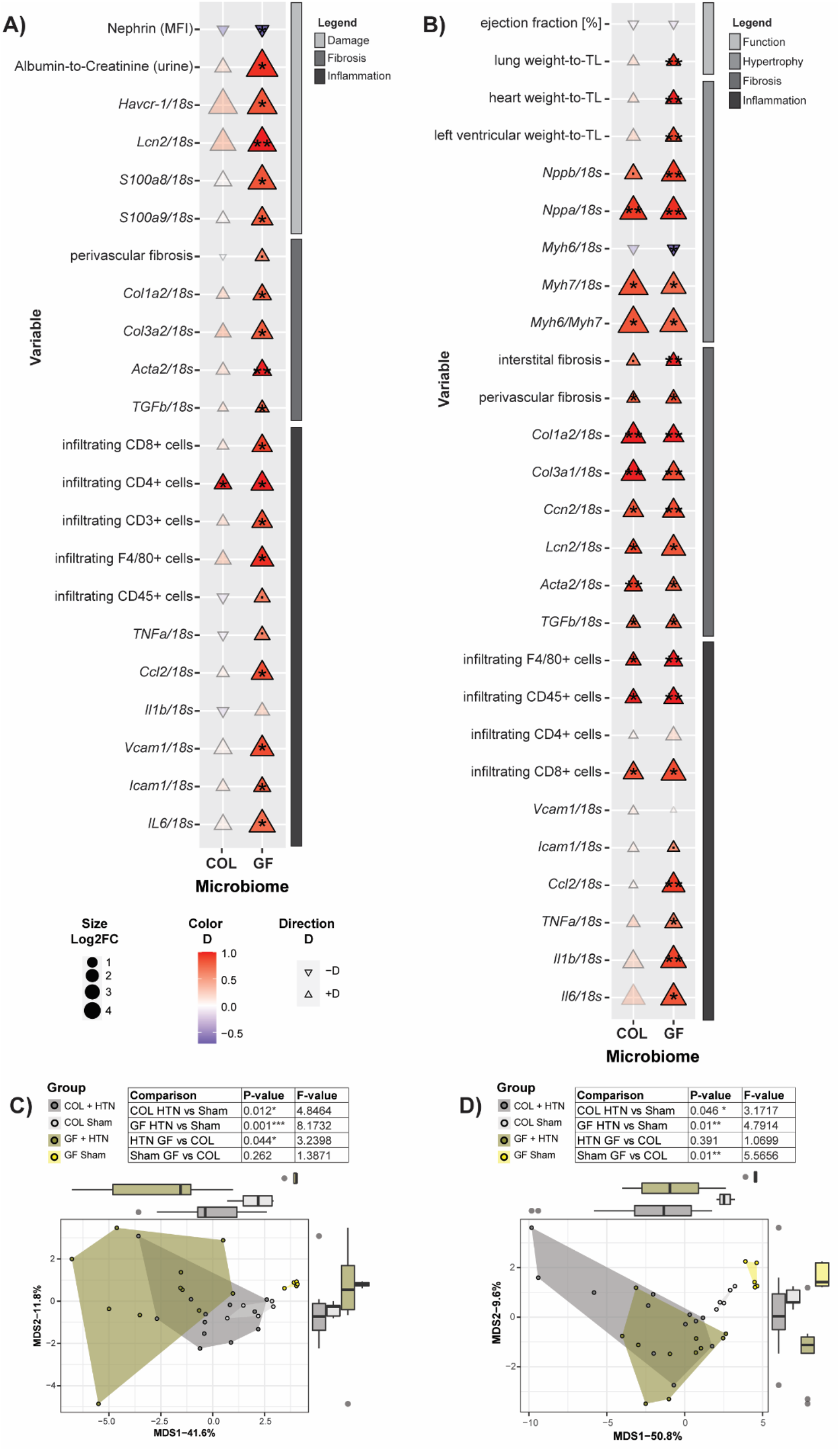
Unbiased assessment of kidney and heart damage in HTN treated GF vs COL mice. A) Effect sizes (Cliff’s delta [D]) and fold changes are shown to assess the effect of HTN treatment relative to the respective sham group in GF and COL on the kidney phenotype. Color and triangle direction indicates effect size. Size indicates log2-transformed fold change (Log2FC) of HTN compared to the respective sham group. Significance was calculated using the Mann-Whitney U-test and was FDR-corrected with the Benjamini-Hochberg procedure to account for multiple testing. Markers for significant Q-values are superimposed, and transparency indicates non-significant effects. °q<0.1, *q<0.05, **q<0.01. Variables are organized by subcategory (Damage, Fibrosis, Inflammation). B) Same univariate analysis as in A) was performed on cardiac phenotypic data. Variables are organized by subcategory (Function, Hypertrophy, Fibrosis, Inflammation). C) Principal Coordinate analysis was performed based on Euclidean distance scaling of kidney phenotypic data to demonstrate the dissimilarities between study groups in a quantitative phenotypic state space. Pairwise comparisons between groups were preformed using PERMANOVA and reported in the inset table. D) Principal Coordinate analysis as in C) was performed on the cardiac phenotypic data.

To further quantify the HTN effect across the renal (Figure 3C) and cardiac (Figure 3D) tissue space, we performed a multivariate principal coordinate analysis (PCoA) summarizing the overall (dis)similarities amongst the groups. To assess pairwise comparisons of interest, the overall dataset was divided and tested using PERMANOVA (Figure 3C). The pairwise comparisons of the kidney phenotypic data show a significant distance in the COL group from sham to HTN (P-value=0.012, F-value= 4.8). However, the effect of HTN, as measured by the F-value, was greater in the GF mice (P-value= 0.001, F-value= 8.2). In line with our initial univariate targeted analysis (Figure 1), this indicates that the overall phenotypic change in the kidney in response to HTN was larger in GF mice than in COL mice. Furthermore, the pairwise PERMANOVA between GF+HTN and COL+HTN groups (P-value= 0.044, F-value= 3.2) indicates that HTN induction resulted in a different outcome on a multivariate level. Interestingly, despite some slight differences within some univariate kidney data in the sham groups (e.g., albuminuria, F4/80+ cells), pairwise comparison of GF and COL sham samples was insignificant (P-value= 0.262, F-value= 1.4).

For the multivariate analysis of our cardiac data, pairwise comparisons were also used to assess the trajectory of each group (Figure 3D). Consistent with our conclusions from the univariate assessment of the overall cardiac phenotype (Figure 3B), we saw that the comparison between GF+HTN and COL+HTN group showed significant overlap in the PCoA plot, and the pairwise comparison between these samples was not significant (P-value= 0.391, F-value=1.1). Conversely, the difference between sham GF and COL samples was significant, suggesting that perhaps the basal cardiac phenotype is sensitive to the host’s microbiome status (P-value = 0.01, F-value = 5.6). As expected, the comparison of HTN to sham samples within the GF (P-value = 0.01, F-value = 4.8) and COL (P-value = 0.046, F-value = 3.2) samples were both significant. The cardiac data again indicate larger phenotypic shifts in GF mice most likely driven by significantly different sham groups.

Taken together, our univariate and multivariate approaches indicate a larger effect of HTN in GF mice. In the case of the kidney, this increased effect is indeed driven by a stronger adverse response of GF mice to HTN, whereas in the case of the heart, this effect is driven by phenotypic differences in the healthy groups (sham-treated mice).

### Vascular response to Angiotensin II is similar in GF and COL mice

There is some evidence from the literature that vascular reactivity may be dependent on microbial colonization.^22, 23^ We opted not to implant telemetry devices for the measurement of BP in our primary experimental animals, as microscopic vascular surgery was not possible under sterile conditions. Although we decided to forgo this gold-standard for BP measurement, we still questioned whether the axenic status of our GF mice would impact the basal mean arterial pressure (MAP), or BP reactivity to Ang II. We therefore colonized additional mice using the same colonization procedure as previously outlined, and we performed *in vivo* BP measurements using an implanted arterial catheter in freely moving mice. Interestingly, we found that GF mice had a significantly higher MAP (Figure 4A) than their colonized counterparts, although the mean of each group (GF mean value = 118.4 mmHg, COL mean value = 107.8 mmHg) was still in a range considered normal for untreated C57BL/6J mice.^24^ Acute intravenous infusion of Ang II induced an increase in BP (Figure 4B) which was nearly identical for GF and COL mice, suggesting that AngII-dependent reactivity of the vasculature is similar in these mice. Similarly, we investigated *ex vivo* vascular contractility of mesenteric arteries isolated from conventionally colonized mice (CONV) or GF mice (GF). CONV mice were used for this *in vitro* experiment due to ease of availability because colonization of GF mice is a lengthy procedure. GF mice showed similar contractile response compared to CONV mice in response to Ang II (Supplemental Figure S5A). Additionally, mesenteric arterial rings from GF mice showed similar contraction force in response to KCl to CONV mice (Supplemental Figure S5B).

**Figure 4.**
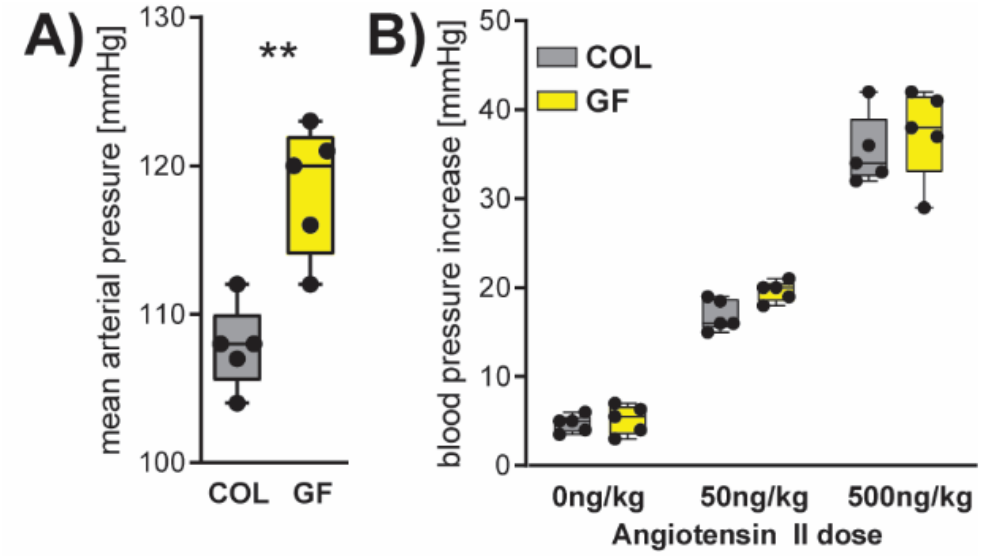
Blood pressure and Angiotensin II reactivity in GF and COL mice. A) GF and COL mice were implanted an arterial catheter for blood pressure measurement. Mean arterial pressure was measured in resting and awake animals, and the difference was tested using an unpaired two-tailed t-test (** P ≤ 0.01). B) The increase in blood pressure after acute intravenous infusion of Angiotensin II as a bolus was measured. No statistical difference between GF and COL mice was found using a two-way repeated measurement ANOVA (p=0.14), whereas Ang II dose had a significant influence (p ≤ 0.0001, not shown).

### Microbiota and microbial metabolites shape serum metabolome changes in hypertension

Although the vascular reactivity of GF mice was similar to those housing microbes, we were interested to understand the different phenotypic and inflammatory response to HTN. We therefore decided to investigate the microbiome itself and associated metabolite production, as our group and others have previously shown that some metabolites of microbial origin can be anti-inflammatory in hypertension.^5, 13, 14^ Consistent with the literature^25^, the metabolome of GF and COL animals was affected by HTN induction. We found a multitude of differences when analyzing the serum metabolome (MxP Quant 500, Biocrates) from GF and COL mice in HTN compared to the respective sham, which we grouped together by class to show the composite shift of over 300 individually measured metabolites (Figure 5A, Supplemental Table 3). Of the 15 classes of metabolites where HTN induction had an effect, four of those classes (phosphatidylcholines, hexosylceramides, alkaloids, fatty acids) showed similar trajectories, suggesting these classes changed in a microbiome-independent manner upon HTN (Figure 5A). Unsurprisingly, the serum metabolome was significantly impacted by the microbiome, and we saw a wide range of correlations between individual metabolites and microbial species, as well as functional modules derived from shotgun sequencing data (Figure 5B). Although the serum metabolite measurements used here were very comprehensive, they did not cover SCFA metabolites, which we and others have shown to have high importance in the progression of HTN. It has been demonstrated elsewhere that GF mice are devoid of some important SCFA^26^. We confirmed this for our mouse colonies (GF and CONV, which were the source populations for our experiments) by performing mass spectrometry measurements of fecal acetate, propionate, and butyrate (Fig 5C). We expect that the lack of the potent anti-hypertensive metabolites propionate and butyrate influences the phenotype seen in GF mice.

**Figure 5.**
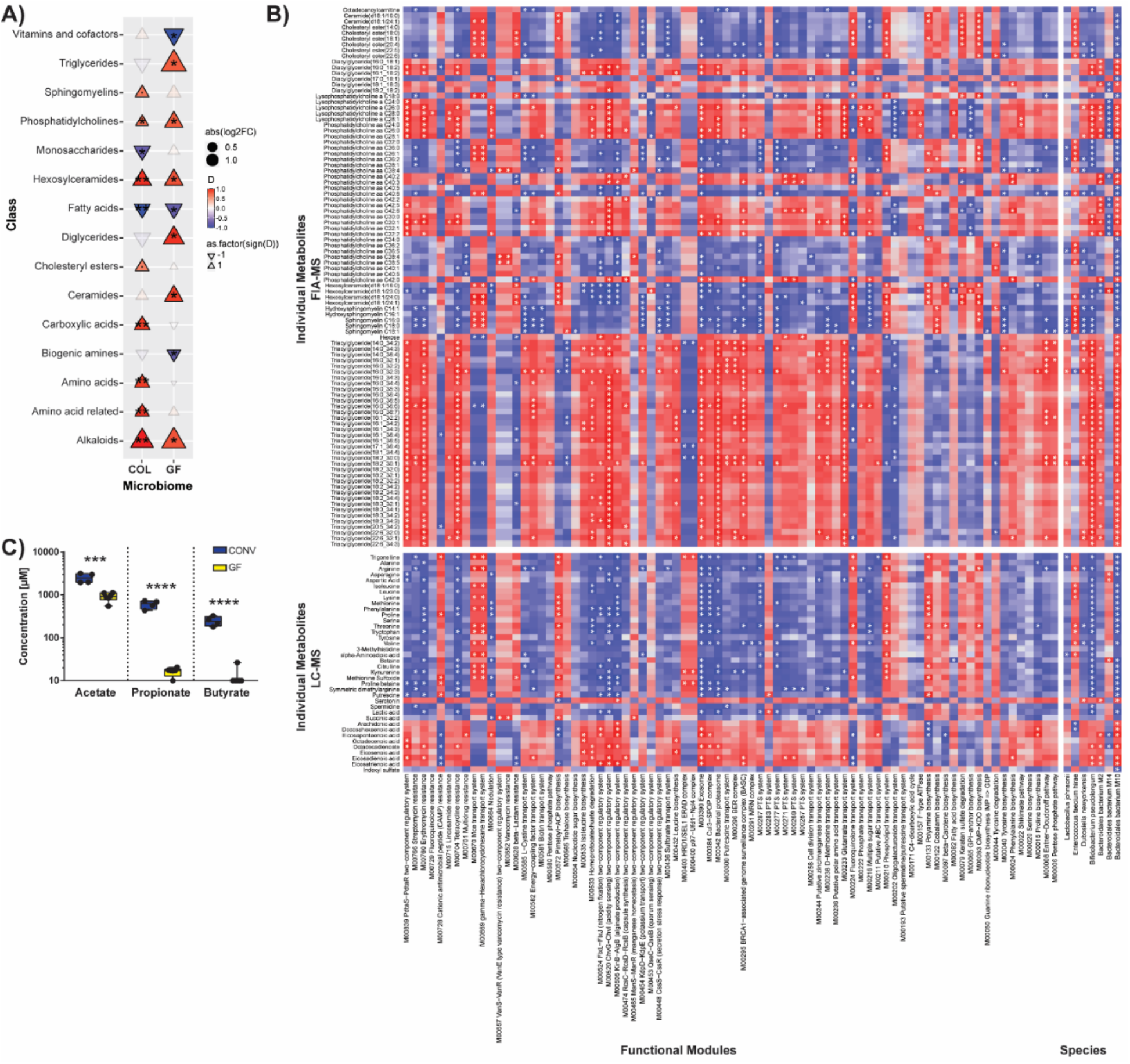
Microbiome heavily influences the metabolome in response to HTN. A) Using the Biocrates MxP500 Quant kit, serum metabolites were analyzed from GF and COL mice. Metabolites were grouped by class to demonstrate the overall effect on the serum metabolome. Effect size (Cliff’s delta [D]) is shown to demonstrate the effect of HTN relative to the respective sham group in GF and COL. Colour and triangle direction indicates effect size. Size indicates log2-transformed fold change (Log2FC) of HTN mice compared to the respective sham group. Significance was calculated using the Mann-Whitney U test and was FDR-corrected using the Benjamini-Hochberg procedure to account for multiple testing. Markers for significant Q-values are superimposed, and transparency indicates non-significant effects. °q<0.1, *q<0.05, **q<0.01. Correlation between individual metabolites, arranged by class, on the y-axis and species, as well as functional microbial modules on the x-axis within the COL group are shown in (B). Variables which were significantly (Mann-Whitney U test) altered between HTN and sham were chosen, Spearman’s correlations were calculated, and FDR corrected for multiple testing (superimposed in white (*q<0.05)). C) Fecal short-chain fatty acids (SCFA) from GF or CONV mice demonstrates the absence of potent anti-hypertensive metabolites butyrate and propionate in the GF setting (GF n=6, CONV n=4). Significance was tested using unpaired two-tailed Student’s t-test (***p ≤ 0.001, ****p ≤ 0.0001). LC-MS, liquid chromatography-mass spectrometry; FIA-MS, flow injection analysis-mass spectrometry.

### Inflammation contributes to the differing phenotypic response to HTN in GF and COL mice

It has been shown that the immune system and the gut microbiome are strongly interconnected, and several studies have shown the importance of immune cells in HTN.^2, 3^ We and others have also shown that metabolites of microbial origin can influence inflammation in HTN.^5, 13, 14^ We examined the splenic immune cell composition a surrogate for systemic inflammation in GF and COL mice by flow cytometry. Of the 23 immune cell subsets we investigated, 12 were differentially influenced by HTN when compared to the respective sham group (Fig 6A, Supplemental Table 4). We then examined all immune parameters multivariately to assess how changes in immune cells would be reflected in the distance between each group using PCoA (Figure 6B). It is known that the immune system of GF mice differs from their colonized counterparts^27^. Our data confirms this observation as the pairwise comparison of sham-treated GF and COL mice (P-value = 0.011, F-value = 6.6) was significant. Interestingly, this difference between COL and GF was still evident after HTN induction (P-value = 0.026, F-value = 2.8). Pairwise comparisons of individual groups using PERMANOVA indicated that the HTN to sham comparison was significant in the GF group (P-value = 0.006, F-value = 4.8), but not within the COL group (P-value = 0.177, F-value = 1.6) (Figure 6B). It can be concluded that overall, the inflammatory status of GF mice was disturbed to a greater degree by HTN induction.

**Figure 6.**
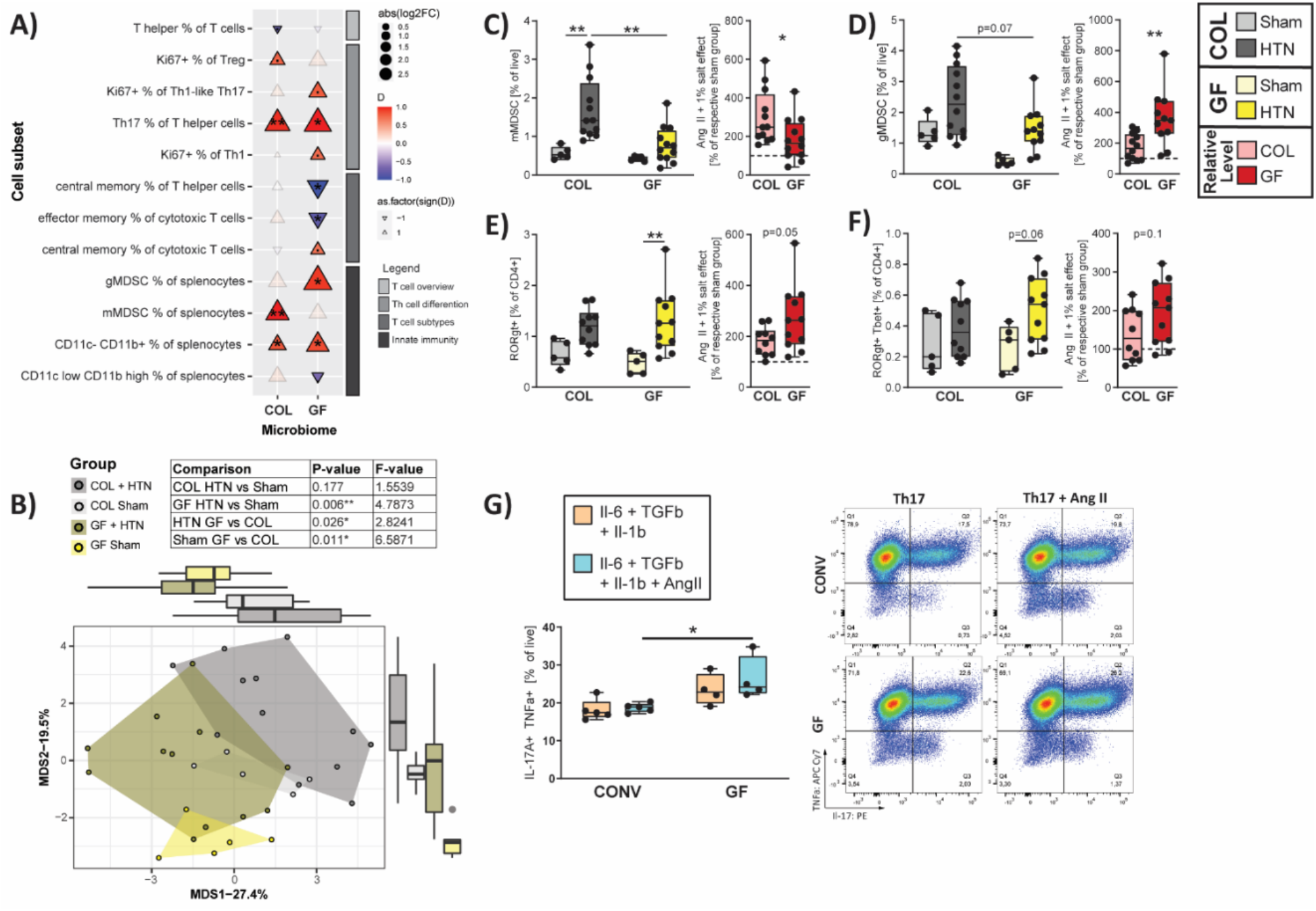
Colonization status influences the inflammatory reaction to HTN. A) Several unique immune cell subsets were measured from the spleen, as a proxy measure of systemic inflammation. Effect size (Cliff’s delta [D]) is shown to demonstrate the effect of HTN relative to the respective sham group in GF and COL. Color and triangle direction indicates effect size. Size indicates log2-transformed fold change (Log2FC) of HTN induction compared to the respective sham group. Significance was calculated using the Mann-Whitney U test and was FDR-corrected using the Benjamini-Hochberg procedure to account for multiple testing. Markers for significant Q-values are superimposed, and transparency indicates non-significant effects. °q<0.1, *q<0.05, **q<0.01. Immune cell subsets are organized by subcategory (Innate immunity and T cell overview, differentiation, and subtypes) and subsets, and cell types where no significant effect in either GF or COL was found are not shown. B) Principal Coordinate analysis was performed based on Euclidean distance scaling of immune cell abundances to demonstrate the dissimilarities between study groups. Pairwise comparisons between groups were performed using PERMANOVA and reported in the inset table. Monocytic MDSCs (mMDSC, C) and granulocytic MDSCs (gMDSC, D), as well as Th17 (E) and Th1-like Th17 cells (F) from the spleen are shown as boxplots on the left. G) Naïve T cells from mesenteric lymph nodes of either GF of CONV mice, polarized *in vitro* towards a Th17 using a cocktail of IL-1β, TGFβ, and IL-6 with or without Ang II (GF n= 4, CONV n= 5). To the left in (C-G), two-way ANOVA and post-hoc Sidak multiple comparison’s test was used to test significance. To the right in (C-F), the relative change induced by Ang II + 1% NaCl in comparison to the respective sham group was tested using an unpaired two-tailed T-test. No change (100%) depicted as dotted line. For all plots in (C-G), p-values are as follows; * P ≤ 0.05, ** P ≤ 0.01.

Previous studies have shown that splenic MDSC increase in HTN and have anti-hypertensive properties.^9^ In line with the literature, there was an increase in monocytic MDSC (mMDSC) upon HTN induction, and this increase was only significant in COL+HTN mice (Figure 6C). Furthermore, for both mMDSC and granulocytic MDSC (gMDSC) (Figure 6D) subtypes, COL+HTN mice showed a significantly higher frequency of these anti-hypertensive immune cells than GF+HTN mice. Interestingly, the relative increase in HTN compared to sham for mMDSC was lesser in GF mice but greater for gMDSC than in COL (Figure 6C-D). The dynamics of whether the relative increase, or absolute number of MDSCs is relevant in HTN is currently unknown. Additionally, we saw an increase in Th17 cells in both GF and COL mice, though this effect only reached significance in GF mice (Figure 6E). This increase was driven by pathological Th1-like Th17 cells, defined by their co-expression of RORgt and Tbet (Figure 6F).^28^ Recent evidence suggest that pre-existing conditions can influence naïve T cell responses towards effector differentiation.^29-31^ We hypothesized that the absence of microbes and their metabolites in GF mice would render naïve T cells more vulnerable to cytokines. Therefore, we performed an *in vitro* Th17 polarization of naïve T cells from GF and CONV mice (which were used in place of COL due to ease of accessibility) in the presence or absence of Ang II. Naïve T cells from GF mice more readily polarized towards Th17 cells than cells from CONV mice, particularly in the presence of Ang II (Figure 6G). To demonstrate that the conditions for the polarization of Th17 cells indeed exist *in vivo*, we quantified the expression of each of the Th17-polarizing cytokines (Il-6, Il-1b, and TGFb) by qPCR from kidney (Supplemental Figure S6A, Figure 3A) and heart (Supplemental Figure S6B, Figure 3C) tissue. These key cytokines were increased upon HTN. The comparison of effect sizes (Figure 3A, C) often indicated larger effects of HTN in GF mice.

In summary, our *in vitro* findings suggest that the different pre-conditioning of naïve T cells within GF and COL mice may impact polarization into pro-inflammatory effector T cells and thus severity of target organ damage. We anticipate that this effect contributes to our *in vivo* findings, where the absence of microbes had a potent impact on immune cells relevant in HTN and cardiorenal damage.

## Discussion

Gut microbial dysbiosis associates with HTN in humans^5, 32^ and in rodent models.^5, 32-34^ Interestingly, fecal microbiota transplantation from hypertensive patients into mice has also been shown to induce an increase in blood pressure.^32^ However, few studies have focused on the overall contribution of the microbiome to the pathogenesis of HTN, such that it may be contextualized relative to other known contributors to hypertensive disease burden. Here we show in a presence/absence scenario, that the microbiota has a potent effect on HTN-induced cardiac and renal damage in mice. GF mice showed a stronger adverse response to HTN than their COL littermates. Interestingly, the kidney seems to be more sensitive to changes in microbial status than the heart. Lastly, we propose that the altered inflammatory response in GF mice contributes to their aggravated phenotype in HTN.

We have shown robustly that the kidney damage within GF mice upon HTN is comparatively more severe than the damage experienced by their COL littermates. Some of the larger effects in our univariate analysis may be related to the slightly lower baseline level of putative markers for kidney damage within sham GF mice compared to sham COL, though these groups do not significantly differ. We surmise that the baseline differences between GF and COL mice are likely due to the immunological uniqueness of GF mice, which has been documented in the literature.^27^ Indeed, across renal damage, fibrosis and inflammation markers, we consistently saw a significant effect of HTN selectively in the GF group (Fig 3A). Furthermore, we show on a multivariate level that the difference between sham GF and GF+HTN mice for the composite of kidney parameters is significant, while the equivalent comparison within the COL mice is insignificant (Figure 3C). Consistent with other inflammatory markers, we show an increase of infiltrating macrophages (F4/80+ cells, Figure 1E) in GF+HTN kidneys, as well as the increased expression of *Ccl2* (Supplemental Figure S3A), which has been implicated as a major player in worsening kidney damage in mice^35^ and humans^36^. Infiltrating macrophages during renal injury are known to contribute to the secretion of cytokines like IL-1β, which enhances the activation and differentiation of Th17 cells.^37^ A previous study additionally showed that SCFA-treatment in ischemia-reperfusion injury (IRI) radically reduced kidney *Ccl2, Il-1b*, and associated kidney damage.^38^ SCFA have been shown to have anti-inflammatory properties in several cell types^39-42^, which could contribute to the lessened damage in COL mice, since only mice harboring microbiota have significant SCFA production within the gastrointestinal tract (Figure 5C). Our results are highly compatible with previous studies showing that GF status exacerbates kidney damage in the context of IRI^43^ and adenine-induced chronic kidney disease^20, 44^.

Intriguingly, we found that the cardiac phenotype was less influenced by the microbial status of the host. Here we have shown that particularly for markers of fibrosis (Figure 3B), regardless of the microbiome status, the mice developed significant injury. For CD8+ T cells, F4/80+ macrophages and overall CD45+ leukocytes, we observed significant changes in GF and COL mice, although GF mice tended to show a higher fold change in response to HTN (Supplemental Table 2). Despite these similarities, both cardiac hypertrophy (Figure 2A) and left ventricular mass-to-tibia length (Figure 2B) were significantly altered upon HTN induction in the GF but not COL mice. Nonetheless, our data suggest that the kidney, more so than the heart, represents a subspace of hypertensive target organ damage, which is more susceptible to microbial colonization. It is conceivable that cardiac damage could be further exacerbated as renal function declines. Thus, the gut microbiota could be added as an important modulator of the well-known cardio-renal axis. Further research to follow up this idea is required, perhaps using several iterations of variations to a defined community of microbes, to test the universality of this hypothesis.

Metabolites of microbial origin, some of which are known to be associated with cardiovascular disease and accumulate in chronic kidney disease^16, 17^, were measurable within the serum metabolome of our COL but not our GF mice, such as IS and TMAO (Supplemental Figure S1D). Our results very clearly indicate that GF mice experience robust kidney damage to a greater extent than COL mice, despite GF mice being devoid of these harmful metabolites. However, we suggest that the reason COL mice experience less overall damage is likely due to the presence of SCFA. We and others have shown the potent effect of SCFA in mouse models.^13, 14^ Here we have shown again, for a representative set of animals, that SCFA are depleted in GF mice (Figure 5C). We hypothesize that the potency of SCFA in COL mice counterbalances the presence of IS and TMAO. Further research on this topic is required to definitively conclude the effects of the co-occurrence of these various metabolites of microbial origin.

Furthermore, we show that systemic inflammatory response to HTN is altered by colonization status. MSDC, which represent an important subset of innate anti-inflammatory cells in HTN^9^, reacted differently GF mice compared to COL (Figure 6C-D). Additionally, we found that Th17 cells were increased during HTN in GF mice (Figure 6E). Th1-like Th17 cells, which are known to be pathogenic, trended towards enrichment in GF+HTN mice (Figure 6F). We wanted to explore *in vitro* that naïve T cells from GF mice were more sensitive to polarizing cytokines and Ang II. We found that upon polarization, naïve T cells from GF mice skewed more towards Th17, particularly when Ang II was added (Figure 6G). As it has been recently demonstrated that the SCFA propionate can decrease the rate of Th17 cell differentiation^41, 42^, we suspect that this could be part of the reason naïve cells from COL mice were less inducible toward Th17. Recently, Krebs and colleagues showed that the development of Th17 cells in the kidney is dependent on the cytokine micromilieu and can be blocked with specific antibodies against IL-1b and IL-6.^45^ We also could show that the polarization conditions used in our Th17 *in vitro* assay were practically available *in vivo*, and the expression of each polarizing cytokine was increased within heart and kidney tissue of hypertensive mice.

To our knowledge, one study similar to ours exists within the literature, published by Karbach et al.^46^ It is clear from the extensive phenotyping performed in our study that our findings were not congruent with their data, where they showed that GF mice were protected from developing HTN and related vascular damage. Though this was initially a surprise, upon further examination, there are two likely scenarios that may explain this. First, the protocol of our experiments did differ from one another. The study from Karbach and colleagues compared GF mice to conventionally raised mice, whereas we compared GF mice with littermates that had been colonized early in life. Therefore, our study was able to account for known genetic drifts in gnotobiotic colonies. Additionally, Ang II infusion was only performed for seven days, while we studied a more chronic phenotype. Furthermore, our mice ate different diets, and as Kaye and colleagues recently demonstrated, the composition of the diet can have a profound impact on the resultant hypertensive phenotype.^47^ Second, it is highly likely that the microbiome used in our study and in the study by Karbach and colleagues may be distinct from one another. Incongruencies like ours have also been found in other contexts. The comparison of microbiome-rich and GF mice in one study showed the amelioration of an IRI of the kidney by the microbiome^43^, where another study demonstrated the opposite effect^48^.

To investigate the second scenario further, we hypothesized that the microbiome background used in any given study might have drastic implications for the study outcome. Our group and others have shown that microbially-produced metabolites have a potent effect on the pathogenesis of HTN.^5, 13, 14, 49^ In reference to that, if the microbiota itself were to change, we expect the circulating metabolites to be likewise altered within the host. Unfortunately, the study from Karbach et al.^46^ did not include any information regarding the microbiome and metabolome of their microbiota-rich mice. Although, we did find a recent study from Cheema and Pluznick^25^ where these data were made available, but their phenotypic data is not reported.^25^ Nevertheless, to test our hypothesis regarding the putative comparability of the colonizing microbiome between studies, and the impact this may have on resultant study outcome, we decided to compare our microbiome and metabolome to the published data from Cheema and Pluznick. To compare the microbiomes from these two studies, we re-annotated our shotgun microbiome sequencing data such that it would be comparable to the 16s rDNA sequencing data from Cheema and Pluznick. The microbiome of colonized mice between the two studies were starkly contrasting in sham and HTN mice (shown as a multivariate PCoA plot derived from genus level information from each of the studies, Supplemental Figure S7A). We surmised that because of the lack of overlapping microbiome signatures within our study and the Pluznick dataset, that the metabolome signal would likewise be dichotomous. We compared the serum metabolome dataset from the two studies by using metabolites which could be measured in both studies from all COL and GF mice. We found that interestingly, there was significantly less distance between the effect of HTN on individual metabolites within the serum metabolome of GF mice from these two studies than in the equivalent COL mice comparison (Supplemental Figure S7B-C). This result suggests that the congruence of the serum metabolome in GF groups within these two datasets is higher than the COL groups. These exploratory data support the idea that the structure of the implanted microbiome has a measurable impact on serum metabolome alterations in response to HTN. Because of our and others’ findings regarding the importance of microbial metabolites in HTN, we believe that this could be a driving factor behind the contradictory phenotypic results of our study compared to the data from Karbach et al.^46^. It is nonetheless critical in future studies for the microbiome to be well-documented and openly accessible to avoid questions regarding the reproducibility of existing studies.

## Conclusion

We have shown that the microbiota has a profound effect on hypertensive disease pathogenesis. Furthermore, we have shown that GF mice, when compared to their colonized littermates, experienced an aggravation of target organ damage, which was more distinct in the kidney than in the heart. Additionally, we demonstrated that the metabolome is influenced significantly by the microbiome used for experimentation, which underscores the need for standardization of experimentation and reporting within the field. The immunophenotype of HTN mice, and in particular, the alteration of MDSC and Th17 cells, which have been previously implicated in HTN, give us some indication of how GF mice may have developed an exacerbated hypertensive phenotype in our study. We propose that the COL mice were protected from damage in comparison to their GF counterparts due to the absence of the potent anti-inflammatory SCFA metabolites under GF conditions.

## Author Contributions

N.W., D.N.M., H.B. designed the study. E.G.A., H.B., G.N., D.T., L.M., A.M., A.R. and N.W. performed animal experiments and analyzed the data. A.F.R, M.T., and M.B. performed *in vivo* BP measurement. T.U.P.B. and U.L. performed the microbiome analysis. E.G.A., R.F.G., S.K. and J.A.K performed and analyzed the metabolomics experiments. S.K.F. and C.Y.C. helped with data analysis and interpretation. E.G.A., N.W., H.B., and D.N.M. wrote the manuscript with input from all authors.

## Acknowledgments

We thank Petra Voss, Ilona Kramer, May-Britt Köhler, Jana Czychi, Ute Gerhardt, Alina Eisenberger, Martin Taube, and Stefanie Schelenz for their excellent technical assistance.

