## Supplemental Figures 1-7, Tables 1-9, Methods for "Quantifying the impact of gut microbiota on inflammation and hypertensive organ damage"

### SUPPLEMENTAL METHODS

Animal Ethics

All experiments performed complied with the German/ European law for animal protection and were approved by the local ethics committee (G0280/13, G0028/21). Mice were maintained on a 12:12 hour day: night cycle with constant access to food and water.

Animal Protocol

Wild-type C57BL/6J mice were bred under axenic conditions in an isolator (Metall+Plastic, Radolfzell-Stahringen, Germany). Until week four, mice in both experimental groups grew up under GF conditions. At four weeks of age male mice were randomized to either remain GF in the isolator or receive passive bacterial colonization (COL). For colonization, mice were introduced in the regular SPF animal facility and placed in cages from healthy wild-type male C57BL/6J mice. Until 12 weeks of age mice received sterilized tap water as drinking water. At 12 weeks of age COL and GF mice received Angiotensin II (Ang II, 1.44 mg/kg/d) by subcutaneous infusion via an osmotic minipump (Alzet) and 1% NaCl (Carl Roth) in the drinking water or sham treatment. Minipumps were implanted under sterile conditions, GF mice were kept sterile throughout the experiment. Sterile drinking water was delivered via the Hydropac system (Plexx B.V., Elst, Netherlands). Throughout the experiment mice were fed autoclaved standard breeding chow (V1124, Ssniff, Soest, Germany). After two weeks of Ang II + 1% NaCl or sham treatment mice were euthanized by isoflurane anesthesia and blood, spot urine (where possible), feces, and organs were collected. As a control group for in vitro experiments, cells, and tissues from age-matched conventionally raised SPF C57BL/6J mice (referred to as CONV) were used.

Echocardiography

Echocardiography was performed as described by Markó et al.[1] Briefly, mice were anesthetized with isoflurane and examined on the VisualSonics Vevo 2100 system using a 30 MHz-Transducer (MS-400, VisualSonics). Left ventricular wall thickness was analyzed using M-mode images from the parasternal short-axis view. All measurements were performed by an experienced reader blinded to treatments.

*In vivo* blood pressure and Ang II pressor response

Acute mean arterial pressure (MAP) was measured in unrestrained conscious germ-free and colonized controls. Basal cardiovascular parameters as well as Ang II pressor response were obtained within 4 hours after exposing germ-free mice to unsterile air. To record blood pressure and infuse Ang II intravenously, self-customized vascular catheters were used.[2] Mice were deeply anesthetized with isoflurane (4-5% for induction; 1-2% for maintenance), and the catheter implantation performed on a thermo controlled surgery table as previously described.[3] Briefly, a small incision over the femoral triangle was made to expose the femoral artery and vein in which heparin/saline filled (100 U/mL) catheters were introduced ~1.5 cm proximally, reaching the abdominal aorta and vena cava, respectively. The catheters were tunneled subcutaneously, exteriorized, and secured between the scapulae with a silk suture 3/0. Before surgery mice received a subcutaneous injection of Carprofen (5mg/Kg) for pain management, and recovered from anesthesia individually in a cage placed on a thermo controlled plate at 37 °C. After consciousness regain, the arterial catheter was connected to a pressure transducer (AD instruments #MLT0699) to record baseline cardiovascular parameters beat-by-beat for approximately ~30 minutes. Data was acquired using a PowerLab data acquisition system (PowerLab/4sp) and a LabChart software v5. Once baseline parameters were recorded, the maximal pressor response to increasing doses of Ang II (0, 50 and 500 ng/kg) were calculated by the difference between the peak MAP response and the averaged immediate (~1 minute before Ang II infusion) baseline MAP. Ang II (Calbiochem #05-23-0101) was injected in bolus, intravenously, connecting a 100 µL Hamilton syringe (Hamilton #80600) to the intravenous catheter. Aliquots of Ang II were prepared to inject the volume of 0.5 µL/g body weight. Each Ang II bolus injection consisted of total final volume of 100 µL, containing the Ang II solution followed by a saline washout. Ang II doses were administered with a ~10 minutes interval in between when minimal physical activity was observed.

Immunophenotyping

Mice were euthanized by isoflurane anesthesia; spleens were removed and kept at 4 °C in PBS with 0.5 % bovine serum albumin (BSA, Sigma Aldrich) and 2 mM EDTA (Sigma Aldrich). Single-cell suspensions were obtained using 70 µm strainers, followed by erythrocyte lysis, and further filtering using a 40 µm mesh. Cells were counted with trypan blue exclusion for flow cytometry analysis and $10^6$ cells were used per multicolor flow cytometry panel. All flow cytometric measurements included dead cell exclusion using Live/Dead Fixable Aqua Dead Cell Stain Kit, for 405 nm excitation (Thermo Fisher). Cells were stained with surface antibodies (Supplemental Table 5) in PBS + 0.5 %BSA + 2 mM EDTA together with Fc blocking reagent (Miltenyi) for 30 min on ice. Intracellular antigens were stained using FoxP3 Staining Buffer Kit (eBioscience). Respective intracellular antibodies (Supplemental Table 5) were incubated for 30 min on ice. Data was recorded on a BD FACS Canto II using BD FACS Diva software. Data analysis was performed with FlowJo (TreeStar).

*In vitro* Th17 polarization

Cells were isolated from mesenteric lymph nodes of GF or conventionally raised C57BL6/J mice. These cells were then sorted using magnetic activation with the CD4+ T Cell Isolation Kit (Miltenyi Biotec) according to the manufacturer's instructions. Isolated CD4+ T cells were collected and resuspended in MACS buffer at 2 x $10^6$ in 50 µL. For APC-free differentiation, cells were fluorescently stained for 30 min in an antibody cocktail containing anti-CD4–PercyP Vio 700, anti-CD44–PE, anti-CD62L–APC and anti-CD25–PE–Cy5, and subsequently purified by fluorescence-activated cell sorting on a BD FACS Aria II Flow Cytometry Cell Sorter (BD Biosciences). Sorted naïve CD4+ T cells (CD4+CD62L+CD44lowCD25−) were stimulated by plate-bound anti-CD3 (2 µg ml−1, 145-2C11, BD Pharmingen) and anti-CD28 (2 µg ml−1, 37.51, BD Pharmingen) in the presence of Th17 polarizing conditions using IL-6 (40 ng $ml^{-1}$), rhTGFβ1 (2 ng $ml^{-1}$) and Il-1b (10 ng $ml^{-1}$). To determine the influence of Ang II on Th17 cell polarization in GF or COL, naïve CD4+ T cells were cultured with vehicle used for stock dilution ($H_20$) or 50 µL Ang II for 96 hrs. Cells were stained and measured as detailed above for Th17-specific markers. Antibodies used are shown in Supplemental Table 5.

Urine analysis

Spot urine was collected from a subset of experimental mice upon sacrifice. All parameters were measured with the AU480 clinical chemistry analyzer (Beckman Coulter) according to the manufacturer's instructions. The measurement of urinary albumin was done by a turbidimetric approach, where absorption of light is proportional to the albumin concentration. The measurement of creatinine was done by an enzymatic conversion of creatinine and colorimetric quantification of the reaction.

Histology staining of renal and cardiac tissue

Heart and renal tissue were immediately frozen in -40 °C isopentane and subsequently stored at -80 °C. Staining was performed on 5 µm cryosections fixed in -20 °C acetone for 10 min. Unspecific binding was blocked with 10 % normal donkey serum (NDS) for 2 hrs.

For cardiac cryosections, primary antibodies were diluted in 10 % NDS (anti-CD4 (H129.19), BD Pharmingen, 1:75; anti-CD8a (53-6.7), BD Pharmingen, 1:100; anti-CD3-A647 (17A2), BioLegend, 1:50; anti-F4/80 (A3-1), Abcam, 1:100; anti-type I collagen (polyclonal), Southern Biotech, 1:100; anti-fibronectin (polyclonal), Abcam, 1:400) and incubated overnight at 4 °C in a humid chamber. Cy3-conjugated secondary antibodies were diluted in PBS for 2 hrs at room temperature (RT) (anti-goat-IgG-Cy3 (polyclonal), Jackson ImmunoResearch, 1:300; anti-rabbit-IgG-Cy3 (polyclonal), Jackson ImmunoResearch, 1:400; anti-rat-IgG-Cy3 (polyclonal), Jackson ImmunoResearch, 1:200) and incubated at room temperature for 2 hrs. Cardiac interstitial fibrosis was analyzed using fibronectin immunofluorescence. Cy-3 positive area was quantified in five high-power fields (HPF, 40x) per heart slide of crosscut cardiomyocytes using ImageJ software with a mean threshold for the Cy3-positive area. Cardiac perivascular fibrosis was quantified using type I collagen immunofluorescence. Cy3-positve fibrosis area was measured using CaseViewer software (3D Histech) and normalized to the mean vessel area. Representative images of interstitial and perivascular fibrosis were taken with an 80x magnification in *Caseviewer*. Infiltrating immune cells were counted using CD4- and CD8-specific antibodies (with associated DAPI-positive nuclei) and quantified per heart section using CaseViewer software (3D Histech). F4/80 and CD3-positive (with DAPI-positive nuclei) cells were counted and quantified in five high-power fields (HPF, 40x) per heart section using CaseViewer software (3D Histech).

For kidney cryosections, primary antibodies were diluted in 10 % NDS (anti-CD4 (H129.19), BD Pharmingen, 1:100; anti-CD8a (53-6.7), BD Pharmingen, 1:100; anti-F4/80 (A3-1), Abcam, 1:100; anti-CD45 (D3F8Q), Cell Signalling, 1:200; anti-CD3-A647 (17A2), BioLegend, 1:50; anti-Nephrin (polyclonal), RD Systems, 1:20) and incubated overnight at 4 °C in a humid chamber. Cy3-conjugated secondary antibodies were diluted in PBS (anti-goat-IgG-Cy3 (polyclonal), Jackson ImmunoResearch, 1:300; anti-rabbit-IgG-Cy3 (polyclonal), Jackson ImmunoResearch, 1:400; anti-rat-IgG-Cy3 (polyclonal), Jackson ImmunoResearch, 1:200) and incubated at room temperature for 1-2 hrs. For the analysis of Nephrin, five evenly distributed cortical regions per sample were analyzed by taking 20x magnification fields of view in the *Caseviewer* software. The further image analysis was performed in *FIJI*. Glomeruli that had their main body in the section plane (approximately round, comparably sized) were chosen for analysis. Of these glomeruli, the mean fluorescence intensity was measured within the area of each glomeruli (determined by encircling the glomeruli by hand). Representative images of one glomerulus per group were taken with an 80x magnification in *Caseviewer* and equally color adjusted in *FIJI*. Infiltrating immune cells were counted as CD4-, CD8-, F4/80-, CD45- and CD3-positive (with DAPI-positive nuclei) cells and quantified in five high-power fields (HPF, 40x) equally dispersed in the cortical region of each kidney section using CaseViewer software (3D Histech).

Kidney tissue was additionally fixed in 4% formalin PBS solution, washed after 24 to 72 hrs in PBS, and then embedded in Paraffin and cut to 3 µm. Formalin fixed tissue was stained with Masson's trichrome using standard protocols. Perivascular fibrosis of the kidney was quantified as described above using Masson's trichrome staining. All histological stainings were scanned using a Pannoramic MIDI II slide scanner (3D Histech).

Quantitative real-time RT-PCR of renal and cardiac tissue

Half a kidney and the heart apex were shock-frozen in liquid nitrogen and stored at -80 °C. RNeasy Mini Kit (QIAGEN) was used for isolation of RNA following the manufacturer's protocol. RNA concentration and quality were determined using a NanoDrop-1000 Spectrophotometer (PeqLab). cDNA was synthesized from 2 µg RNA using the Applied Biosystems High-Capacity cDNA Reverse Transcription Kit (Thermo Fisher). TaqMan or SYBR Green assays were used to quantify target gene expression using the standard curve method on a QuantStudio 3 (Thermo Fisher). Target mRNA expression was normalized to the *18S* housekeeping gene. All primers and probes were designed with PrimerExpress 3.0 (Applied Biosystems) and synthetized by Biotez (Berlin, Germany). All primer and probe sequences are provided in Supplemental Table 6.

Wire-myography in isolated mesenteric arteries

First-order mouse mesenteric arteries were removed immediately after sacrificing the animals under inhalation anesthesia with isoflurane by cervical dislocation, quickly transferred to cold (4°C), oxygenated (95% $O_2$/5% $CO_2$) physiological salt solution (PSS) containing (in mmol/L) 119 NaCl, 4.7 KCl, 1.2 $KH_2PO_4$, 25 $NaHCO_3$, 1.2 $MgSO_4$, 11.1 glucose, 1.6 CaCl2. The vessels were dissected into 2 mm (mesenteric artery) rings whereby perivascular fat and connective tissue were removed without damaging the adventitia. Mesenteric rings were positioned on two stainless steel wires (diameter 0.0394mm) in a 10-ml organ bath of a Mulvany Small Vessel Myograph (DMT 610 M; Danish Myo Technology, Denmark). The organ bath was filled with 5 ml PSS. The bath solution was continuously oxygenated with a gas mixture of 95% $O_2$ and 5% $CO_2$ and kept at 37 °C (pH 7.4). The mesenteric rings were placed under a tension equivalent to that generated at 0.9 times the diameter of the vessel at 100 mm Hg by stepwise distending vessels using LabChart DMT Normalization module. This normalization procedure was performed to obtain the passive diameter of the vessel at 100 mm Hg. The software Chart5 (AD Instruments Ltd. Spechbach, Germany) was used for data acquisition and display. After 60 min equilibration, vessels were pre-contracted with phenylephrine (60 mM KCl) until a stable resting tension was acquired. The composition of 60 mM KCl (in mmol L) was 63.7 NaCl, 60 KCl, 1.2 $KH_2PO_4$, 25 $NaHCO_3$, 1.2 $Mg_2SO_4$, 11.1 glucose and 1.6 $CaCl_2$. Ang II was directly added to the bath solution. Tension is expressed as a percentage of the steady-state tension (100%) obtained with KCl. Salts and other chemicals were obtained from Sigma-Aldrich (Germany).

Metabolomics

*Materials* - All reagents, internal and calibration standards, quality controls, and a patented 96-well filter plate required for MxP Quant 500 analysis were included in the kit provided by Biocrates Life Science AG (Innsbruck, Austria).

*MxP Quant 500 assay and sample preparation* - The MxP Quant 500 kit from Biocrates Life Science AG is a kit-based assay based on phenylisothiocyanate (PITC) derivatization of the target analytes using internal standards for quantification. Up to 630 metabolites from 26 analytical classes can be measured including small molecules (1 alkaloid, 1 amine oxide, 20 amino acids, 30 amino acid related, 4 bile acids, 9 biogenic amines, 1 carbohydrate, 7 carboxylic acids, 1 cresol, 12 fatty acids, 4 hormones, 4 indoles and derivatives, 2 nucleobases and related and 1 vitamin) and lipids (40 acylcarnitines, 76 phosphatidylcholines, 14 lysophosphatidylcholines, 15 sphingomyelins, 28 ceramides, 8 dihydroceramides, 19 hexosylceramides, 9 dihexosylceramides, 6 trihexosylceramides, 22 cholesteryl esters, 44 diglycerides and 242 triglycerides).

Plate preparation was done according to the manufacturer's protocol. Briefly, 10 µL of serum was transferred to the upper 96-well plate and dried under a nitrogen stream. Thereafter, 50 µL of a 5% PITC solution was added. After incubation, the filter spots were dried again before the metabolites were extracted using 5 mM ammonium acetate in methanol (300 µL) into the lower 96-well plate for analysis after further dilution using water. Internal standards were present in the plate prior to analysis. Calibration standards at different dilutions (Cal 1 to Cal 7) were also included on the same plate according to Biocrates recommend layout. Biocrates quality control (QC) samples were run every 20 samples. One pooled QC sample was measured at the beginning, two in the middle and one at the end.

*MxP Quant 500 measurement*- Evaluation of the instrument performance prior to sample analysis was assessed by the Biocrates recommended system suitability test (SST). Separate test mixtures were provided with the kit for liquid chromatography-mass spectrometry (LC-MS) and flow injection analysis-mass spectrometry (FIA-MS) SST evaluation. Pass/Fail criteria were according to Biocrates recommended criteria.

The LC-MS system was comprised of a 1290 Infinity UHPLC-system (Agilent, Santa Clara, CA, USA) coupled to a 5500 QTrap with SelexIon (AB Sciex Germany GmbH, Darmstadt, Germany) in electrospray ionization (ESI) mode. Acquisition method parameters are shown in Supplemental Table 7-9. Two LC methods and two FIA methods were used to analyze the same set of samples. Raw data was assessed and manually curated in Met*IDQ* version Nitrogen (Biocrates Life Science AG, Innsbruck, Austria). Compounds were identified and quantified using isotopically-labeled internal standards and pre-determined multiple reaction monitoring (MRM) transitions for LC and by their accurate mass for FIA, as according to the Biocrates protocol.

*Data analysis* - The Met*IDQ* software is provided by Biocrates and is usually used for data processing and normalization. Normalization was done on median of Biocrates QC level 2 in 4 replicates. The concentrations of metabolites that were analyzed by FIA were automatically calculated by the software based on internal standard ratios. The analyte peaks obtained by LC were integrated by the Sciex Analyst version 1.6.3 software and normalized to the internal standards. The absolute concentrations for 42 compounds were determined based on the 7-point calibration curve. All quantification was done within MetIDQ version Nitrogen. An in-house developed script was used for data quality analysis and preprocessing which was a modified version of Metaquac[2]. The modification enabled the retention of metabolites considered above the limit of detection but below the limit of quantification by Biocrates software. The median relative standard deviation (RSD) for pooled QC samples was 8.35 % for FIA and 8.81 % for LC.

*Final metabolite identification* - The metabolites were considered valid when they appeared in a minimum of 70% of biological replicates. Only analytes with values above the limit of detection (LOD) were considered. The LOD for individual analytes was defined as three times the median peak area in the blank samples (peak intensity was used for FIA data) and a minimum intensity of 1 000 count per second (cps). Analytes below the LOD were rejected. Each analyte was subsequently normalized to its respective labeled internal standard. Data was quality checked among other things for technical reproducibility, missing values and batch effects, as detailed previously.[4] Outliers were removed where a drastically low number (less than 70% of the overall average metabolite count) of metabolites were capturable (likely an issue occurred during sample preparation), or where there was consistently greater than 2 standard deviations between a given sample and the mean for its respective biological group. The filtered and quality checked dataset was used for subsequent statistical analysis. Where necessary, imputation of group-specific mean value for individual metabolites was used, although this was rare as 98.7% and 99.1% of data points were available in the final selection of GF and COL data, respectively.

*Fecal SCFA measurement* - Short chain fatty acids from C57BL6/J GF and CONV mouse fecal matter were isolated as described in Haghikia et al.[5] Extracted samples were measured using GC-MS performed on a Thermo Scientific™ Q Exactive™ hybrid quadrupole Orbitrap mass spectrometer. Data analysis was preformed using Thermo Scientific™ Xcalibur™ Software.

Microbiome analysis

*DNA Isolation and Extraction* - All laboratory procedures were conducted under a hood with laminar flow (LabGarda ES Energy Sever Classe II Laminar Flow, NuAire Inc., Plymouth, Minnesota) to limit environmental contamination. Total DNA was extracted from all stool samples using the ZymoBIOMICS DNA Miniprep Kit (ZYMO Research Europe GmbH, Freiburg, Germany). Using 20-50 mg aliquots from the previously frozen stool and caecum content, DNA extraction was performed by following the manufacturer's recommendations with slight modifications for better mechanical disruption. The samples were added to the PeQLab vial containing the ZR BeashingBead Lysis Tube beads (0.1 & 0.5 mm), and after adding 750 µL ZymoBIOMICS Lysis Solution, the vial was caped tightly. Samples were mechanically disrupted using a PeQLab Precellys 24 (Bertin Corp., Rockville, USA) for 2 x 15 s at 5500 RPM. After 5 min rest, the cycle was repeated. The remaining steps of the DNA extraction procedure followed the manufacturer's protocol by ZYMO Research Europe GmbH, the final elution of DNA was performed with 100 µL ZymoBIOMICS DNase/RNase Free Water. The samples were stored at -20° C before being shipped on dry ice to Novogene for sequencing.

The extraction protocols were also performed on a positive control (stool samples from CONV mice) and with a blank sample (RNase-, and DNase-free, genomic DNA-free water). A 75 µL blank and 20-50mg positive control was used for extraction.

*Microbiome Sequencing and analysis* - Shotgun sequencing data were processed using ngless.[6] Quality control was performed by trimming reads with phred score below 25, while trimmed reads shorter than 45 bp were discarded. Taxonomic classification was assessed by aligning the quality trimmed reads to mOTUs[7] (v2.5) using bwa[8] (v0.7.17) with default parameters.

Mouse genome GRCm38.p6 was masked with the sequences retrieved from SILVA[9] database V138 using bbtools (https://sourceforge.net/projects/bbmap/). The reads which passed quality control were mapped to the masked mouse reference using bwa within ngless. Sequences matching the mouse genome were considered off-targets and discarded. The microbial composition among different samples and within groups of samples was visualized using KronaTools (v2.7).[10]

Trimmed and filtered reads were aligned to the global microbial gene catalogue (gmgc v1.0), covering 2.3 billion ORFs from 13,174 metagenomes and 14 habitats, as described previously.[11] Reads which mapped to multiple genes in the reference database were handled using the "dist1" method and normalized using the "scaled" mode. The results were parsed by a customized perl script and visualized using the ggplot2[12] package. Functional annotation was performed mapping the trimmed and filtered reads to the MouseGutCatalog (v0.9)[13], while the dist1 method was used to account for multiple annotations.

*Fecal qPCR* - For all stool and caecum content samples and all relevant controls, PCR was performed using an Applied Biosystems QuantStudio 3 system (Thermo Fisher Scientific, Darmstadt, Germany). Amplification and detection were performed in 96-well optical plates (Applied Biosystems) with SYBR-Green (Applied Biosystems). For amplification, the standard protocol of the Applied Biosystems QuantStudio 3 system was followed, i.e., an initial cycle at 95 °C for 10 min, followed by 40 cycles at 95 °C for 15 s, and 1 min at 60 °C. To check for specificity, melting curve (Tm) analysis was performed, increasing the temperature from 60 to 95 °C at a rate of 0.2 °C per second with the continuous monitoring of fluorescence.

Standard curves for quantification consisted of ten-fold serial dilutions in the range of $10^8$ –$10^0$ copies of the 16S rRNA gene of the E. coli (Invitrogen, C404010) amplified with primers 27F (5'-GTTTGATCCTGGCTCAG-3') and 1492R (5'-CGGCTACCTTGTTACGAC-3'). Using universal primers, Univ 337F 5'-ACTCCTACGGGAGGCAGCAGT-3' and Univ 518R 5'-GTATTACCGCGGCTGCTGGCAC-3' we quantified the total amount of bacterial 16S in stool and caecum content samples (COL and CONV), in stool from GF mice, as well as in the blank and positive controls. Copy numbers per g feces was calculated for each primer set used, as previously described.[14, 15]

Statistical Analyses

All statistical analyses were performed using either R version 4.0.2 or GraphPad Prism 6. From the COL+HTN group n=1 animal was removed from the cardiac RNA data due to technical failure of the RNA preparation. For the collagen I staining n=1 GF sham mice was removed due to technical failure of the staining. Univariate data was analyzed using two-way ANOVA, where the interrogated factors were HTN and microbiome status. When one or both factors were significant, post-hoc testing with Sidak's multiple comparison test was performed to identify the source of variation. For the assessment of relative changes, values from HTN mice were expressed in percent of the mean of the respective sham group. COL and GF were then compared using unpaired two-tailed Student's t-test. $p<0.05$ was considered statistically significant. Multivariate principal coordinate analysis and subsequent testing using permutational multivariate analysis of variance (PERMANOVA) was performed in R using the vegan[16] package, and Euclidean distances (kidney, heart, metabolome and immunome data) or Bray-Curtis (microbiome data) were used for dissimilarity indices. For the heatmaps, COL+HTN and GF+HTN mice were compared to their respective sham group using Mann-Whitney U test. Multiple comparisons were adjusted using Benjamini-Hochberg correction for false discover rate. $q<0.1$ was considered statistically significant. Effect size analysis was performed using Cliff's delta in R using orddom[17] and visualized with ggplot2[12] packages. Additional in-depth description of microbiome and metabolome statistical analysis and graphical approaches are detailed in below.

*Metabolite-Microbiome comparison*- Metabolites and microbes on genus level were first subjected to filtration, where only features that have significantly changed between sham and HTN based on Mann-Whitney U test were selected. Spearman's correlation between selected features were calculated and followed by FDR correction on the P-values. Same procedure was followed to obtain the Spearman's correlation between low-concentration metabolites and functional modules. These two matrices were combined and the heatmap was plotted by using the R package gplots.[18]

*Comparative analysis with Pluznick study data: Microbiome*- Raw 16S sequences of fecal microbiome sequencing of Pluznick study were downloaded from NCBI Sequence Read Archive (BioProject: PRJNA514044). Then LotuS 1.62[19] was used for analyzing the 16S amplicons based on SILVA[9], Greengenes[20] and HITdb databases, and resulted in all taxonomic levels of microbiome abundance tables. Subsequently, these abundance tables were rarefied by using RTK.[21] Abundance table on genus level was adopted for the comparison with this study. Principal Coordinates Analysis (PCoA) was done with vegan package[16], and Bray-Curtis distances was used.

*Comparative analysis with Pluznick study data: Metabolome*- Metabolites from Pluznick study were curated for overlapping metabolites where there were exact matches for the biochemical name, CAS ID, HMDB ID, and/or PubChem IDs. Metabolite data from the Pluznick study included both male and female mice. Multivariate tests including PERMANOVA and PCoA were performed by using vegan[16] R package, and Euclidean distance was adopted. Comparison of the changes in metabolites between sham and HTN groups on individual level were done by first performing Mann-Whitney U test within GF or COL groups in each study (NAs were removed if any NA resulted from the test). We then compared the directionality of change for each metabolite in the equivalent group from the Pluznick or Berlin data. For the comparison of effect size for each metabolite individually between sham and HTN groups in either the GF or COL/Conv conditions of the Pluznick or Berlin study, Cliff's delta was calculated by using orddom R package.[17] The distances between the effect sizes in GF or COL groups were calculated by taking the absolute value of numerical difference between the two studies. Ggplot2[12] was used for plotting, and ggpubr[22] was applied to add Mann-Whitney U test result.

### SUPPLEMENTAL FIGURES

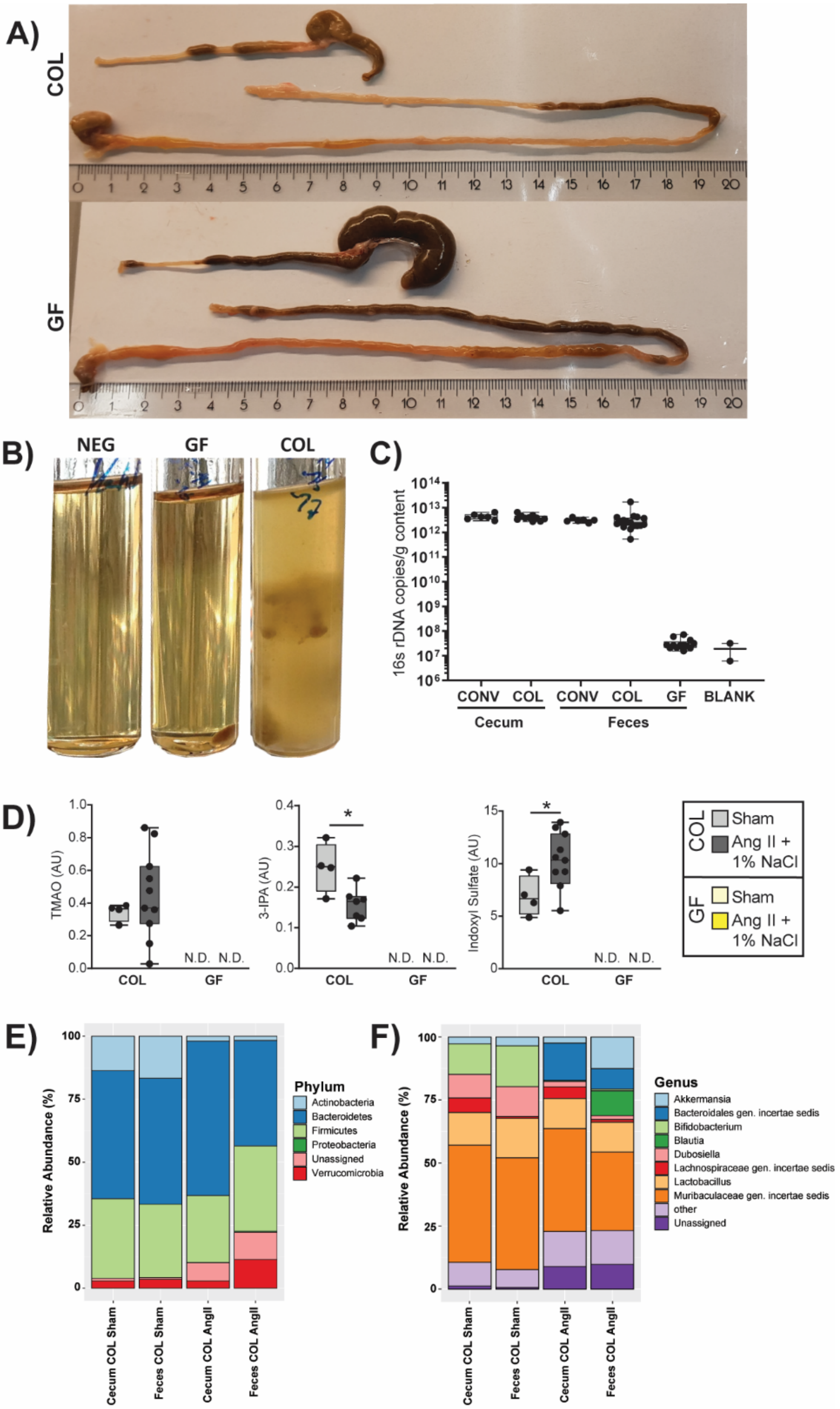

**Supplemental Figure S1.** A) Representative images of GF and COL mouse gastrointestinal tract from stomach to anus upon sacrifice. B) Representative images of thioglycolate bacterial cultivation from fecal pellets GF and COL mice upon sacrifice, compared to a negative control. C) 16s rDNA copies per gram stool from fecal (CONV n= 6, COL n= 17, GF n=13) and cecal content (CONV n= 6, COL n= 10) demonstrates the absence of bacterial DNA in GF mice above background (Blind n= 2). D) Microbially-produced metabolites which were only measurable in the serum of COL and not GF mice using the Biocrates MxP500 Quant kit, including three of notable importance in HTN; TMAO, 3-IPA, and IS. COL group values were tested using an unpaired two-tailed T-test, and * $P \leq 0.05$. Relative abundance of phyla in cecal and fecal matter from COL mice are shown in (E), and genus with greater than 5% relative abundance in at least one subgroup are shown in (F).

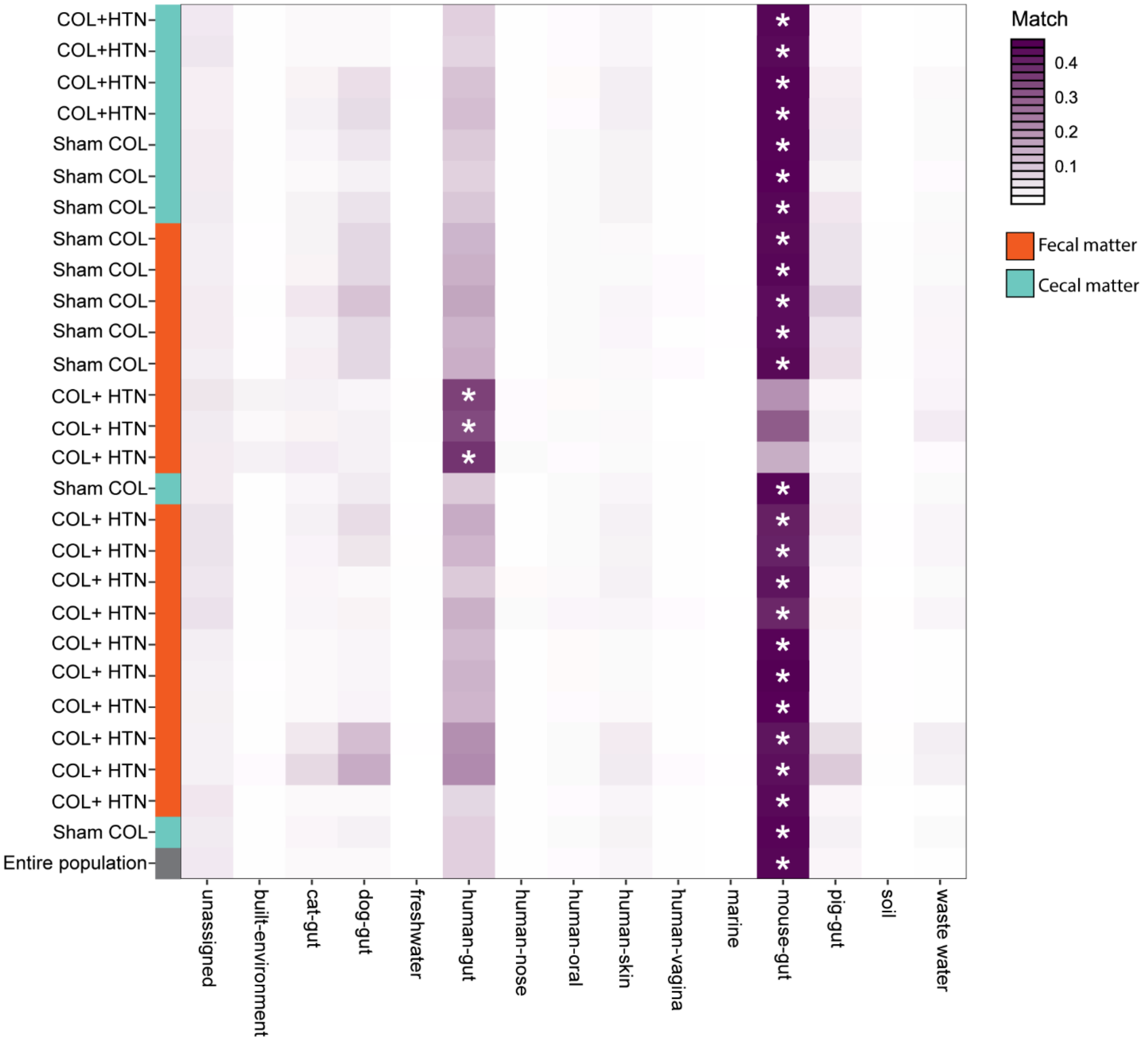


**Supplemental Figure S2.** Annotation of fecal and cecal samples from COL mice to the mouse gut gene catalogue (MGC), indicating the similarity of the mouse microbiome used for experimentation to other communities predefined within this database. * indicates the closest matching environment for a given sample.

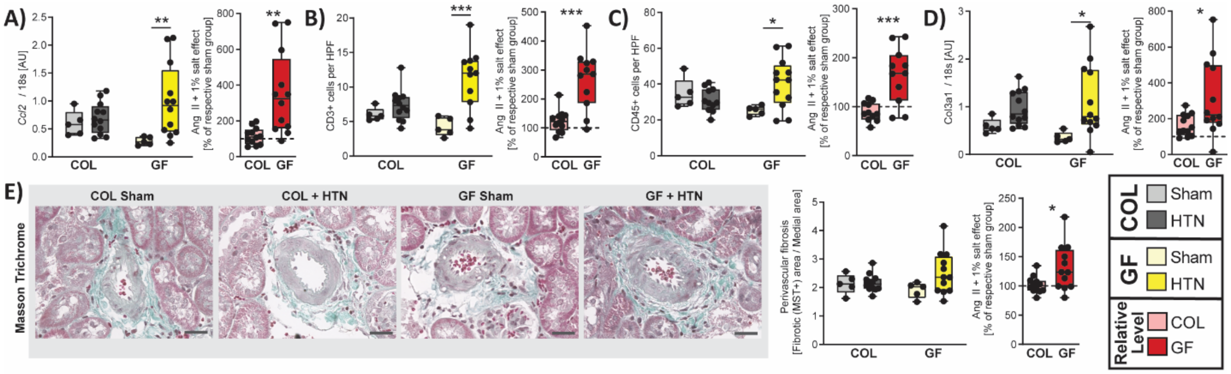


**Supplemental Figure S3.** *Ccl2* (A) and *Col3a1* (D) gene expression measured by qPCR from kidney tissue. CD3+ T cells (B) and CD45+ immune cells (C) were counted from histological images of the kidney. E) Perivascular fibrosis from Masson's Trichrome Staining histological evaluation, measured as the fibrosis positive area divided by the medial area of the vessel. For (A-E), the left graph was tested using a two-way ANOVA and post-hoc Sidak multiple comparison's test. In (A), (B) and (D), hypertension was identified as the source of variation using two-way ANOVA, and post-hoc multiple comparison between sham and HTN within each group revealed that the GF comparison was the source of variation. In the right plot for panels (A-E), the relative change induced by AngII + 1% NaCl in comparison to the respective sham group was tested using an unpaired two-tailed T-test. No change (100%) depicted as dotted line. For all plots, p-values are as follows; * P ≤ 0.05, ** P ≤ 0.01, *** P ≤ 0.001.

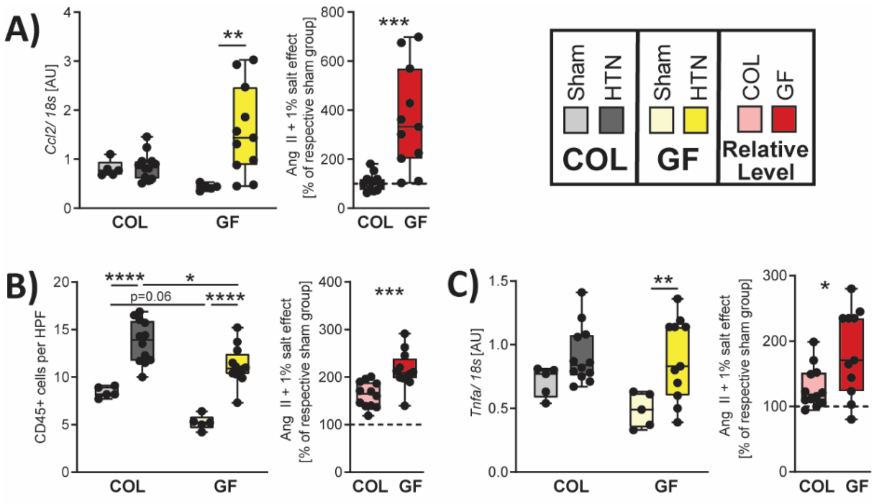


**Supplemental Figure S4.** *Ccl2* (A) and *Tnfa (C)* were measured by qPCR from heart tissue. CD45+ immune cells (B) were counted from 5 representative high-power fields within immunofluorescence stainings of the heart. On the left (A-C), two-way ANOVA and post-hoc Sidak multiple comparison's test was used. In (A) and (C), hypertension was identified as the source of variation using two-way ANOVA, and post-hoc multiple comparison between sham and HTN within each group revealed that the GF comparison was the source of variation. In (B), hypertension and the microbiome were both identified as sources of variation using two-way ANOVA, and all significant post-hoc comparisons are indicated. For (A-C) on the right, an unpaired two-tailed T-test was used to assess the relative change induced by AngII + 1% NaCl in comparison to the respective sham group. No change (100%) depicted as dotted line. For all plots, p-values are as follows; * P ≤ 0.05, ** P ≤ 0.01, *** P ≤ 0.001, **** P ≤ 0.0001.

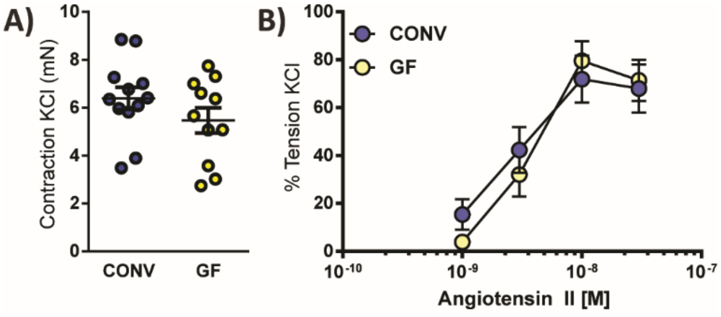


**Supplemental Figure S5.** Mesenteric arteries were isolated from GF and CONV mice. A) Maximal contraction force in response to KCl in mN. B) Contraction of mesenteric arteries in response to increasing doses of Angiotensin II was tested. Contraction force is shown as percentage of the maximal KCl-induced contraction. No statistical difference was found using Student's T-test (A) and two-way ANOVA (B).

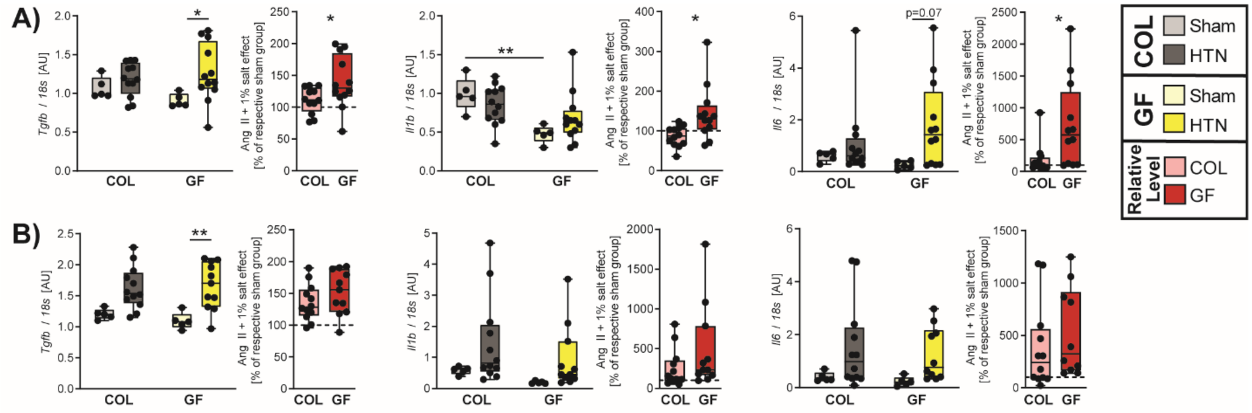


**Supplemental Figure S6.** Kidney (A) and heart (B) expression of cytokine-encoding genes *Tgfb*, *Il1b*, and *Il6*, which are known to induce Th17 cell polarization, measured by qPCR. On the left for each individual marker in (A) and (B), two-way ANOVA and post-hoc Sidak multiple comparison's test was used to test the raw data. For all markers, except kidney *Il1b*, hypertension was identified as the source of variation using two-way ANOVA, and any significant post-hoc multiple comparisons between sham and HTN within each group are shown. For kidney *Il1b*, the source of variation was the microbiome, and post-hoc testing revealed significance stemming from the Sham COL to GF comparison. For each marker, an unpaired two-tailed T-test was used to assess the relative change induced by AngII + 1% NaCl in comparison to the respective sham group, depicted to the right of each raw data plot. No change (100%) depicted as dotted line. For all plots, p-values are as follows; * P ≤ 0.05, ** P ≤ 0.01.

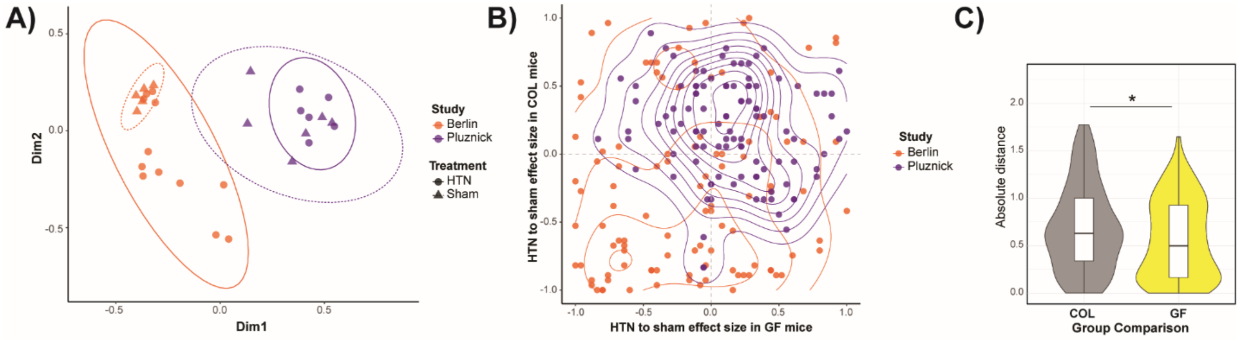


**Supplemental Figure S7.** A) Principal Coordinate analysis was performed based on Euclidean distance scaling of microbiome data published by Cheema and Pluznick (annotated as Pluznick) and our data (annotated as Berlin), to demonstrate the dissimilarities between colonized mice from each study. Colour indicates study group, and ellipse indicates treatment HTN or Sham treatment group. B) Metabolites which were measurable from the serum of mice from GF and COL groups in both the Pluznick and Berlin study, are shown where the effect size of HTN compared to Sham in COL mice is shown on the Y axis, and the effect size of HTN compared to Sham in GF mice is shown on the X axis. Quantification of the distance between the same metabolite on along the GF (X-axis) or COL (Y-axis) from the Pluznick study compared to the Berlin study is shown in (C). *$p<0.05$ by Mann-Whitney U test.

### SUPPLEMENTAL TABLES

#### Table S1.

Univariate analysis from the kidney data space.

| category | variable | P_GF_test | P_COL_test | Q_GF_test | Q_COL_test | FC_GF | FC_COL | log2FC_GF | log2FC_COL | D_GF | D_COL |
|---|---|---|---|---|---|---|---|---|---|---|---|
| Inflammation | *Il6* | 0.019 | 0.646 | 0.023 | 0.782 | 7.349 | 1.871 | 2.878 | 0.904 | 0.733 | 0.167 |
| Inflammation | *Il1b* | 0.104 | 0.383 | 0.104 | 0.660 | 1.444 | 0.862 | 0.530 | -0.214 | 0.533 | -0.300 |
| Inflammation | *Ccl2* | 0.004 | 0.646 | 0.011 | 0.782 | 3.394 | 1.208 | 1.763 | 0.273 | 0.867 | 0.167 |
| Inflammation | *Tnfa* | 0.048 | 0.646 | 0.053 | 0.782 | 1.545 | 0.924 | 0.627 | -0.115 | 0.633 | -0.167 |
| Inflammation | *Icam1* | 0.006 | 0.442 | 0.012 | 0.678 | 1.786 | 1.304 | 0.837 | 0.383 | 0.833 | 0.267 |
| Inflammation | *Vcam1* | 0.004 | 0.721 | 0.011 | 0.830 | 4.485 | 2.017 | 2.165 | 1.012 | 0.867 | 0.133 |
| Fibrosis | *Tgfb1* | 0.014 | 0.279 | 0.020 | 0.583 | 1.341 | 1.101 | 0.423 | 0.139 | 0.767 | 0.367 |
| Fibrosis | *Acta2* | 0.000 | 0.279 | 0.004 | 0.583 | 2.180 | 1.318 | 1.125 | 0.398 | 1.000 | 0.367 |
| Fibrosis | *Col3a2* | 0.009 | 0.027 | 0.014 | 0.154 | 3.165 | 1.638 | 1.662 | 0.712 | 0.800 | 0.700 |
| Fibrosis | *Col1a2* | 0.009 | 0.160 | 0.014 | 0.460 | 2.073 | 1.268 | 1.051 | 0.343 | 0.800 | 0.467 |
| Fibrosis | Perivascular fibrosis | 0.078 | 0.959 | 0.082 | 0.959 | 1.360 | 1.017 | 0.444 | 0.024 | 0.625 | -0.033 |
| Damage | *S100a9* | 0.009 | 0.879 | 0.014 | 0.919 | 3.157 | 1.307 | 1.659 | 0.386 | 0.800 | 0.067 |
| Damage | *S100a8* | 0.006 | 0.879 | 0.012 | 0.919 | 11.424 | 1.880 | 3.514 | 0.911 | 0.833 | 0.067 |
| Damage | *Lcn2* | 0.000 | 0.027 | 0.004 | 0.154 | 11.693 | 6.699 | 3.548 | 2.744 | 1.000 | 0.700 |
| Damage | *Havcr-1* | 0.006 | 0.037 | 0.012 | 0.168 | 16.151 | 12.507 | 4.014 | 3.645 | 0.833 | 0.667 |
| Damage | Urinary albumin-to-creatine ratio | 0.018 | 0.402 | 0.023 | 0.660 | 25.212 | 1.663 | 4.656 | 0.734 | 0.958 | 0.360 |
| Inflammation | Infiltrating CD3+ cells | 0.004 | 0.225 | 0.011 | 0.575 | 2.794 | 1.228 | 1.482 | 0.296 | 0.867 | 0.400 |
| Inflammation | Infiltrating CD4+ cells | 0.002 | 0.002 | 0.011 | 0.021 | 3.715 | 1.875 | 1.894 | 0.907 | 1.000 | 1.000 |
| Inflammation | Infiltrating CD8+ cells | 0.004 | 0.338 | 0.011 | 0.648 | 2.713 | 1.161 | 1.440 | 0.215 | 0.917 | 0.317 |
| Inflammation | Infiltrating CD45+ cells | 0.045 | 0.506 | 0.052 | 0.727 | 1.606 | 0.862 | 0.683 | -0.214 | 0.650 | -0.233 |

| | | | | | | | | | | | |
|---|---|---|---|---|---|---|---|---|---|---|---|
| Inflammation | Infiltrating F4/80+ cells | 0.003 | 0.064 | 0.011 | 0.244 | 4.276 | 1.638 | 2.096 | 0.712 | 0.967 | 0.600 |
| Damage | Nephrin MFI | 0.019 | 0.000 | 0.023 | 0.007 | 0.672 | 0.547 | -0.573 | -0.871 | ##### | -1.000 |

**Table S2.**

Univariate analysis from the heart data space.

| Category | variable | P_GF_test | P_COL_test | Q_GF_test | Q_COL_test | FC_GF | FC_COL | log2FC_GF | log2FC_COL | D_GF | D_COL |
|---|---|---|---|---|---|---|---|---|---|---|---|
| Inflammation | *Il6* | 0.006 | 0.082 | 0.010 | 0.123 | 5.219 | 3.834 | 2.384 | 1.939 | 0.833 | 0.567 |
| Inflammation | *Il1b* | 0.002 | 0.160 | 0.006 | 0.216 | 5.278 | 2.546 | 2.400 | 1.348 | 0.900 | 0.467 |
| Inflammation | *Tnfa* | 0.037 | 0.082 | 0.043 | 0.123 | 1.760 | 1.289 | 0.815 | 0.366 | 0.667 | 0.567 |
| Inflammation | *Ccl2* | 0.001 | 0.721 | 0.006 | 0.743 | 3.492 | 1.064 | 1.804 | 0.090 | 0.933 | 0.133 |
| Inflammation | *Icam1* | 0.082 | 0.574 | 0.092 | 0.646 | 1.212 | 1.140 | 0.278 | 0.189 | 0.567 | 0.200 |
| Inflammation | *Vcam1* | 0.646 | 0.506 | 0.646 | 0.594 | 1.038 | 1.106 | 0.054 | 0.145 | 0.167 | 0.233 |
| Inflammation | Infiltrating CD8+ cells | 0.007 | 0.008 | 0.011 | 0.019 | 4.614 | 2.389 | 2.206 | 1.256 | 0.867 | 0.840 |
| Inflammation | Infiltrating CD4+ cells | 0.205 | 0.743 | 0.221 | 0.743 | 1.493 | 1.071 | 0.578 | 0.098 | 0.417 | 0.127 |
| Fibrosis | *Tgfb1* | 0.006 | 0.009 | 0.010 | 0.019 | 1.508 | 1.352 | 0.592 | 0.435 | 0.833 | 0.800 |
| Fibrosis | *Acta2* | 0.009 | 0.001 | 0.013 | 0.009 | 1.548 | 1.861 | 0.631 | 0.896 | 0.800 | 0.933 |
| Fibrosis | *Lcn2* | 0.006 | 0.002 | 0.010 | 0.010 | 4.110 | 1.648 | 2.039 | 0.721 | 0.833 | 0.900 |
| Fibrosis | *Ccn2* | 0.004 | 0.006 | 0.008 | 0.017 | 2.963 | 2.488 | 1.567 | 1.315 | 0.867 | 0.833 |
| Fibrosis | *Col3a1* | 0.004 | 0.000 | 0.008 | 0.004 | 2.993 | 4.267 | 1.582 | 2.093 | 0.867 | 1.000 |
| Fibrosis | *Col1a2* | 0.000 | 0.000 | 0.004 | 0.004 | 2.507 | 4.252 | 1.326 | 2.088 | 1.000 | 1.000 |
| Fibrosis | Perivascular fibrosis | 0.008 | 0.009 | 0.012 | 0.019 | 1.484 | 1.278 | 0.569 | 0.354 | 0.875 | 0.818 |
| Fibrosis | Interstitial fibrosis | 0.000 | 0.038 | 0.004 | 0.068 | 1.444 | 1.324 | 0.530 | 0.405 | 1.000 | 0.673 |
| Hypertrophy | *Myh7* | 0.019 | 0.006 | 0.025 | 0.017 | 4.731 | 10.249 | 2.242 | 3.357 | 0.733 | 0.833 |
| Hypertrophy | *Myh6* | 0.037 | 0.082 | 0.043 | 0.123 | 0.825 | 0.878 | -0.278 | -0.188 | -0.667 | -0.567 |
| Hypertrophy | *Nppa* | 0.001 | 0.001 | 0.004 | 0.009 | 5.478 | 6.622 | 2.454 | 2.727 | 0.967 | 0.933 |
| Hypertrophy | *Nppb* | 0.001 | 0.027 | 0.006 | 0.052 | 4.192 | 1.770 | 2.068 | 0.823 | 0.933 | 0.700 |

| | | | | | | | | | | | |
|---|---|---|---|---|---|---|---|---|---|---|---|
| Hypertrophy | Left ventricular weight-to-tibia length ratio | 0.002 | 0.160 | 0.006 | 0.216 | 1.604 | 1.314 | 0.682 | 0.394 | 0.900 | 0.467 |
| Hypertrophy | Heart weight-to-tibia length ratio | 0.000 | 0.234 | 0.004 | 0.288 | 1.556 | 1.141 | 0.638 | 0.190 | 1.000 | 0.400 |
| Functional | Lung weight-to-tibia length ratio | 0.004 | 0.221 | 0.008 | 0.284 | 1.295 | 1.175 | 0.373 | 0.232 | 0.960 | 0.418 |
| Functional | Ejection fraction | 0.442 | 0.721 | 0.459 | 0.743 | 0.923 | 0.900 | -0.116 | -0.153 | -0.267 | -0.133 |
| Inflammation | Infiltrating CD45+ cells | 0.002 | 0.002 | 0.006 | 0.010 | 2.146 | 1.629 | 1.101 | 0.704 | 1.000 | 1.000 |
| Inflammation | Infiltrating F4/80+ cells | 0.002 | 0.003 | 0.006 | 0.010 | 1.918 | 1.469 | 0.940 | 0.555 | 1.000 | 0.967 |
| Hypertrophy | *Mhy7/Mhy6* ratio | 0.014 | 0.006 | 0.018 | 0.017 | 7.127 | 13.086 | 2.833 | 3.710 | 0.767 | 0.833 |

**Table S3.**

Univariate analysis from the metabolomics data space.

| variable | P_GF_test | P_COL_test | Q_GF_test | Q_COL_test | FC_GF | FC_COL | log2FC_GF | log2FC_COL | D_GF | D_COL |
|---|---|---|---|---|---|---|---|---|---|---|
| Acylcarnitines | 0.768 | 0.583 | 0.858 | 0.693 | 1.022 | 1.047 | 0.031 | 0.066 | -0.120 | 0.200 |
| Alkaloids | 0.008 | 0.000 | 0.027 | 0.004 | 2.418 | 2.156 | 1.274 | 1.108 | 0.840 | 1.000 |
| Amino acid related | 0.165 | 0.001 | 0.284 | 0.004 | 1.081 | 1.242 | 0.113 | 0.312 | 0.480 | 0.964 |
| Amino acids | 0.679 | 0.002 | 0.858 | 0.006 | 0.988 | 1.306 | -0.018 | 0.385 | -0.160 | 0.927 |
| Bile acids | 0.768 | 0.913 | 0.858 | 0.964 | 9.867 | 1.021 | 3.303 | 0.030 | 0.120 | -0.055 |
| Biogenic amines | 0.013 | 0.221 | 0.027 | 0.300 | 0.831 | 0.875 | -0.267 | -0.192 | -0.800 | -0.418 |
| Carboxylic acids | 0.859 | 0.002 | 0.907 | 0.006 | 0.960 | 1.337 | -0.058 | 0.419 | -0.080 | 0.927 |
| Ceramides | 0.003 | 0.145 | 0.017 | 0.230 | 1.560 | 1.151 | 0.642 | 0.203 | 0.920 | 0.491 |
| Cholesteryl esters | 0.679 | 0.052 | 0.858 | 0.098 | 0.966 | 1.215 | -0.050 | 0.281 | 0.160 | 0.636 |
| Diglycerides | 0.001 | 0.115 | 0.013 | 0.199 | 1.884 | 0.697 | 0.914 | -0.520 | 0.960 | -0.527 |
| Fatty acids | 0.013 | 0.000 | 0.027 | 0.004 | 0.706 | 0.735 | -0.501 | -0.445 | -0.800 | -1.000 |
| Hexosylceramides | 0.008 | 0.001 | 0.027 | 0.004 | 1.571 | 1.570 | 0.651 | 0.651 | 0.840 | 0.964 |
| Lysophosphatidylcholines | 1.000 | 1.000 | 1.000 | 1.000 | 0.962 | 1.014 | -0.056 | 0.020 | 0.000 | 0.018 |
| Monosaccharides | 0.513 | 0.013 | 0.813 | 0.036 | 1.124 | 0.715 | 0.169 | -0.485 | 0.240 | -0.782 |
| Nucleobases and related | 0.679 | 0.913 | 0.858 | 0.964 | 0.940 | 1.103 | -0.090 | 0.142 | -0.160 | 0.055 |
| Phosphatidylcholines | 0.013 | 0.019 | 0.027 | 0.046 | 1.551 | 1.162 | 0.633 | 0.217 | 0.800 | 0.745 |
| Sphingomyelins | 0.075 | 0.038 | 0.143 | 0.080 | 1.235 | 1.218 | 0.305 | 0.284 | 0.600 | 0.673 |
| Triglycerides | 0.013 | 0.267 | 0.027 | 0.339 | 2.649 | 0.719 | 1.405 | -0.476 | 0.800 | -0.382 |
| Vitamins and cofactors | 0.001 | 0.221 | 0.013 | 0.300 | 0.599 | 1.173 | -0.740 | 0.230 | -1.000 | 0.418 |

**Table S4.**

Univariate analysis from the immune data space.

| Category | variable | P_GF_test | P_COL_test | Q_GF_test | Q_COL_test | FC_GF | FC_COL | log2FC_GF | log2FC_COL | D_GF | D_COL |
|---|---|---|---|---|---|---|---|---|---|---|---|
| Th cell differentiation | Ki67+ % of Treg | 0.058 | 0.014 | 0.110 | 0.062 | 1.742 | 1.423 | 0.801 | 0.509 | 0.617 | 0.767 |
| Th cell differentiation | Th17 % of T helper cells | 0.002 | 0.001 | 0.017 | 0.007 | 5.138 | 3.797 | 2.361 | 1.925 | 1.000 | 0.967 |
| Th cell differentiation | Ki67+ % of Th17 | 0.646 | 0.140 | 0.708 | 0.268 | 1.008 | 0.833 | 0.011 | -0.264 | -0.167 | -0.483 |
| Th cell differentiation | Th1-like Th17 % T helper cells | 0.058 | 0.101 | 0.110 | 0.239 | 2.842 | 2.937 | 1.507 | 1.555 | 0.617 | 0.533 |
| Th cell differentiation | Ki67+ % of Th1-like Th17 | 0.013 | 0.104 | 0.050 | 0.239 | 1.729 | 1.287 | 0.790 | 0.364 | 0.800 | 0.533 |
| Th cell differentiation | Th1 % of T helper cells | 0.721 | 0.506 | 0.754 | 0.646 | 0.942 | 0.770 | -0.087 | -0.378 | -0.133 | -0.233 |
| Th cell differentiation | Ki67+ % of Th1 | 0.023 | 0.562 | 0.076 | 0.660 | 1.365 | 1.051 | 0.449 | 0.072 | 0.733 | 0.200 |
| T cell overview | T cells % of splenocytes | 0.874 | 0.342 | 0.874 | 0.492 | 1.001 | 0.963 | 0.002 | -0.055 | -0.067 | -0.317 |
| Th cell differentiation | Treg % of T helper cells | 0.442 | 0.574 | 0.508 | 0.660 | 0.876 | 0.818 | -0.191 | -0.289 | -0.267 | -0.200 |
| T cell overview | T helper % of T cells | 0.082 | 0.013 | 0.145 | 0.062 | 0.942 | 0.931 | -0.086 | -0.104 | -0.567 | -0.800 |
| T cell subtype | Central memory % of T helper cells | 0.002 | 0.916 | 0.017 | 0.916 | 0.440 | 1.134 | -1.184 | 0.181 | -0.983 | 0.050 |
| T cell subtype | Effector memory % of T helper cells | 0.246 | 0.646 | 0.333 | 0.675 | 1.159 | 1.097 | 0.212 | 0.134 | 0.383 | 0.167 |
| T cell subtype | Naive % of T helper cells | 0.130 | 0.234 | 0.213 | 0.385 | 1.069 | 0.915 | 0.096 | -0.128 | 0.500 | -0.400 |
| T cell subtype | Effector memory % of cytotoxic T cells | 0.006 | 0.130 | 0.028 | 0.268 | 0.578 | 1.359 | -0.791 | 0.442 | -0.833 | 0.500 |
| T cell subtype | Central memory % of cytotoxic T cells | 0.031 | 0.460 | 0.088 | 0.622 | 1.333 | 0.928 | 0.415 | -0.108 | 0.700 | -0.250 |
| T cell subtype | Naive % of cytotoxic T cells | 0.225 | 0.646 | 0.324 | 0.675 | 1.034 | 0.953 | 0.048 | -0.070 | 0.400 | -0.167 |
| T cell overview | gd T cells % of T cells | 0.315 | 0.315 | 0.382 | 0.484 | 0.951 | 1.067 | -0.073 | 0.093 | -0.333 | 0.333 |
| Innate immunity | gMDSC % of splenocytes | 0.001 | 0.082 | 0.017 | 0.235 | 5.865 | 1.755 | 2.552 | 0.812 | 0.933 | 0.567 |
| Innate immunity | mMDSC % of splenocytes | 0.160 | 0.000 | 0.245 | 0.007 | 1.684 | 3.000 | 0.752 | 1.585 | 0.467 | 1.000 |

| | | | | | | | | | | | |
|---|---|---|---|---|---|---|---|---|---|---|---|
| Innate immunity | CD11c low CD11b high % of splenocytes | 0.037 | 0.027 | 0.093 | 0.103 | 0.852 | 1.667 | -0.231 | 0.738 | -0.667 | 0.717 |
| Innate immunity | CD11c high CD11b+ % of splenocytes | 0.057 | 0.186 | 0.110 | 0.329 | 0.732 | 0.755 | -0.450 | -0.405 | -0.617 | -0.433 |
| Innate immunity | CD11c+ CD11b- % of splenocytes | 0.292 | 0.082 | 0.373 | 0.235 | 0.881 | 1.170 | -0.182 | 0.227 | -0.350 | 0.567 |
| Innate immunity | CD11c- CD11b+ % of splenocytes | 0.004 | 0.006 | 0.022 | 0.047 | 2.795 | 1.677 | 1.483 | 0.745 | 0.867 | 0.833 |

**Table S5.**
Antibodies used for flow cytometry.

| antigen | fluorophore | clone | concentration | manufacturer |
|---|---|---|---|---|
| CD3ε | VioBlue | 17A2 | 1:10 | Miltenyi |
| CD4 | APC Vio770 | GK1.5 | 1:10 | Miltenyi |
| CD8a | PerCP Cy5.5 | 53-6.7 | 1:100 | eBioscience |
| CD11b | PE | M1/70 | 1:150 | BD Bioscience |
| CD11c | PerCP Cy5.5 | N418 | 1:100 | BioLegend |
| CD25 | VioBright FITC | 7D4 | 1:10 | Miltenyi |
| CD44 | FITC | IM7 | 1:100 | BD Bioscience |
| CD45R / B220 | Alexa Fluor 647 | RA3-6B2 | 1:100 | BioLegend |
| CD62L | APC | MEL-14 | 1:100 | BD Bioscience |
| CD69 | PE Cy7 | H1.2F3 | 1:100 | BD Bioscience |
| γδ-TCR | PE | GL3 | 1:100 | BD Bioscience |
| F4/80 | Pacific Blue | BM8 | 1:100 | BioLegend |
| FoxP3 | PerCP Cy5.5 | FJK-16s | 1:50 | eBioscience |
| Gr-1 | PE Cy7 | RB6-8C5 | 1:150 | eBioscience |
| Helios | Pacific Blue | 22F6 | 1:50 | BioLegend |
| Ki67 | PE Vio770 | REA183 | 1:10 | Miltenyi |
| Ly6C | APC eFluor 780 | HK1.4 | 1:10 | eBioscience |
| RORγt | APC | REA278 | 1:10 | Miltenyi |
| Tbet | PE | REA102 | 1:10 | Miltenyi |
| CD4 | Percyp Vio 700 | GK1.5 | 1:20 | Miltenyi |
| CD44 | PE | IM7.8.1 | 1:20 | Miltenyi |
| CD62L | APC | MEL-14 | 1:100 | BioLegend |
| CD25 | PE–Vio770 | PC61 | 1:100 | BD Pharmingen |
| Il-17A | PE | eBio17B7 | 1:50 | eBioscience |
| TNFa | APC Cy7 | MP6-XT22 | 1:50 | BioLegend |

**Table S6.**

PCR primer and probe information. For: forward primer, rev: reverse primer, probe: Taqman probe.

| gene | | Sequences (5’→3’) |
|---|---|---|
| *18s* | for | ACA TCC AAG GAA GGC AGC AG |
| | rev | TTT TCG TCA CTA CCT CCC CG |
| | probe | FAM – CGC GCA AAT TAC CCA CTC CCG AC – TAMRA |
| *Havcr-1* | for | CTG GAG TAA TCA CAC TGA AGC AAT C |
| | rev | GAT GCC AAC ATA GAA GCC CTT AGT |
| | probe | FAM – CTC CAG GGA AGC CGC AGA AAA ACC – TAMRA |
| *Lcn2* | for | TGA TCC CTG CCC CAT CTC T |
| | rev | GGA ACT GAT CGC TCC GGA A |
| | probe | FAM – TCA CTG TCC CCC TGC AGC CAG A – TAMRA |
| *S100a8* | for | TCA CCA TGC CCT CTA CAA GA |
| | rev | CCA ATT CTC TGA ACA AGT TTT CG |
| *S100a9* | for | TCA GAC AAA TGG TGG AAG CA |
| | rev | GTC CAG GTC CTC CAT GAT GT |
| *Col1a2* | for | CTA CTG GTG AAA CCT GCA TCC A |
| | rev | GGG CGC GGC TGT ATG AG |
| | probe | FAM – CCC ACC CTG TAA ACA CCC CAG CGA AG – TAMRA |
| *Col3a1* | for | CTC ACC CTT CTT CAT CCC ACT CTT A |
| | rev | ACA TGG TTC TGG CTT CCA GAC AT |
| *Tgfb1* | for | CCC GAA GCG GAC TAC TAT GC |
| | rev | TAG ATG GCG TTG TTG CGG T |
| *Vcam1* | for | CTA CAA GTC TAC ATC TCT CCC AGG AA |
| | rev | CAC AGC ACC ACC CTC TTG AA |

| | | |
|---|---|---|
| | probe | FAM – ACA ACG ATC TCT GTA CAT CCC TCC ACA AGG – TAMRA |
| *Icam1* | for | CAG TCC GCT GTG CTT TGA GA |
| | rev | CGG AAA CGA ATA CAC GGT GAT |
| | probe | FAM – CTG TGG CAC CGT GCA GTC GTC C – TAMRA |
| *Tnfa* | for | CGT CCC CAA AGG GAT GAG AA |
| | rev | TGA GGG TCT GGG CCA TAG AA |
| | probe | FAM – TTC CCA AAT GGC CTC CCT CTC ATC A – TAMRA |
| *Ccl2* | for | GGC TCA GCC AGA TGC AGT TAA |
| | rev | CCT ACT CAT TGG GAT CAT CTT GCT |
| | probe | FAM – CCC CAC TCA CCT GCT GCT ACT CAT TCA – TAMRA |
| *Il1b* | for | CGT GGA CCT TCC AGG ATG AG |
| | rev | GAG GAT GGG CTC TTC TTC AAA G |
| *Il6* | for | GTT GCC TTC TTG GGA CTG ATG |
| | rev | GGG AGT GGT ATC CTC TGT GAA GTC T |
| | probe | FAM – TGG TGA CAA CCA CGG CCT TCC C – TAMRA |
| *Fn1* | for | GGA CCT GCA AAC CTA TAG CTG AGA |
| | rev | CTC CCC CAC GAC GTA GGA |
| | probe | FAM – TGT TTT GAT CAT GCT GCT GGG – TAMRA |
| *Nppb* | for | GAA AGT CTC CAG AGC AAT TCA |
| | rev | GGG CCA TTT CCT CCG ACT T |
| *Nppa* | for | AGG AGA AGA TGC CGG TAG AAG A |
| | rev | GCT TCC TCA GTC TGC TCA CTC A |
| | probe | FAM – AGG TCA TGC CCC CGC AGG C – TAMRA |
| *Myh6* | for | GCC AAG ACT GTC CGG AAT GA |
| | rev | TGG AAG ATC ACC CGG GAC TT |

| | | |
|---|---|---|
| *Myh7* | for | CAA TGC CAG GAT TGA GGA TGA |
| | rev | CGT GCC TGA AGC TCC TTG AG |
| *Ccn2* | for | CAA CCG CAA GAT CGG AGT GT |
| | rev | CAC CGA CCC ACC GAA GAC |
| | probe | FAM – CAC TGC CAA AGA TGG TGC ACC CTG – TAMRA |
| *Acta2* | for | TCC TGA CGC TGA AGT ATC CGA TA |
| | rev | GGT GCC AGA TCT TTT CCA TGT C |
| | probe | FAM – AAC ACG GCA TCA TCA CCA ACT GGG A – TAMRA |

**Table S7.**
Ultra-high performance liquid chromatography (UHPLC) gradient for LC part.

| | LC1 | | | | LC2 | | | |
|---|---|---|---|---|---|---|---|---|
| No | Time [min] | Flow [mL/min] | A [%] | B [%] | Time [min] | Flow [mL/min] | A [%] | B [%] |
| 1 | 0.00 | 0.8 | 100 | 0 | 0.00 | 0.8 | 100 | 0 |
| 2 | 0.25 | 0.8 | 100 | 0 | 0.25 | 0.8 | 100 | 0 |
| 3 | 1.50 | 0.8 | 88 | 12 | 0.50 | 0.8 | 75 | 25 |
| 4 | 2.70 | 0.8 | 82.5 | 17.5 | 2.00 | 0.8 | 50 | 50 |
| 5 | 4.00 | 0.8 | 50 | 50 | 3.00 | 0.8 | 25 | 75 |
| 6 | 4.50 | 0.8 | 0 | 100 | 3.50 | 0.8 | 0 | 100 |
| 7 | 4.70 | 1.0 | 0 | 100 | 4.70 | 1.0 | 0 | 100 |
| 8 | 5.00 | 1.0 | 0 | 100 | 5.00 | 1.0 | 0 | 100 |
| 9 | 5.10 | 1.0 | 100 | 0 | 5.10 | 1.0 | 100 | 0 |
| 10 | 5.80 | 1.0 | 100 | 0 | 5.80 | 1.0 | 100 | 0 |

**Table S8.**
Ultra-high performance liquid chromatography (UHPLC) gradient for flow injection (FIA) part.

| No | Time [min] | Flow [mL/min] | A [%] | B [%] |
|---|---|---|---|---|
| 1 | 0.0 | 0.03 | 0 | 100 |
| 2 | 1.6 | 0.03 | 0 | 100 |
| 3 | 2.4 | 0.20 | 0 | 100 |
| 4 | 2.8 | 0.20 | 0 | 100 |
| 5 | 3.0 | 0.03 | 0 | 100 |

**Table S9.**

Mass spectrometry acquisition method parameters for liquid chromatography (LC) and flow injection analysis (FIA). MS: mass spectrometry.

| Option | Parameter | LC1 | LC2 | FIA1 | FIA2 |
|---|---|---|---|---|---|
| MS | Scan type | MRM | MRM | MRM | MRM |
| | Polarity | Positive | Negative | Positive | Negative |
| | MRM detection window (sec) | 30 | 30 | - | - |
| | Duration (min) | 5.45 | 5.45 | 2.95 | 2.95 |
| | Delay time (sec) | 0 | 0 | 0 | 0 |
| | Cycle (sec) | 0.25 | 0.15 | N/A | N/A |
| Advanced MS | Resolution Q1 | Unit | Unit | Unit | Unit |
| | Resolution Q3 | Unit | Unit | Unit | Unit |
| | Intensity threshold | 0 | 0 | 0 | 0 |
| | Setting time (ms) | 0 | 0 | 0 | 0 |
| | Pause between mass ranges (ms) | 2 | 2 | 5.007 | 3 |
| Source/ gas | Curtain gas | 45 | 20 | 20 | 10 |
| | Collision gas | 9 | 8 | 9 | 9 |
| | Ion spray voltage | 5500 | -4500 | 5500 | 5500 |
| | Temperature | 500 | 650 | 200 | 350 |
| | Ion source gas 1 | 60 | 40 | 40 | 30 |
| | Ion source gas 2 | 70 | 40 | 50 | 80 |

6. Coelho LP, Alves R, Monteiro P, Huerta-Cepas J, Freitas AT, Bork P. NG-meta-profiler: fast processing of metagenomes using NGLess, a domain-specific language. Microbiome 2019;**7**(1):84.
7. Milanese A, Mende DR, Paoli L, Salazar G, Ruscheweyh HJ, Cuenca M, Hingamp P, Alves R, Costea PI, Coelho LP, Schmidt TSB, Almeida A, Mitchell AL, Finn RD, Huerta-Cepas J, Bork P, Zeller G, Sunagawa S. Microbial abundance, activity and population genomic profiling with mOTUs2. Nat Commun 2019;**10**(1):1014.
8. Li H, Durbin R. Fast and accurate short read alignment with Burrows-Wheeler transform. Bioinformatics 2009;**25**(14):1754-60.
9. Quast C, Pruesse E, Yilmaz P, Gerken J, Schweer T, Yarza P, Peplies J, Glockner FO. The SILVA ribosomal RNA gene database project: improved data processing and web-based tools. Nucleic Acids Res 2013;**41**(Database issue):D590-6.
10. Ondov BD, Bergman NH, Phillippy AM. Interactive metagenomic visualization in a Web browser. BMC Bioinformatics 2011;**12**:385.
11. Mende DR, Letunic I, Maistrenko OM, Schmidt TSB, Milanese A, Paoli L, Hernandez-Plaza A, Orakov AN, Forslund SK, Sunagawa S, Zeller G, Huerta-Cepas J, Coelho LP, Bork P. proGenomes2: an improved database for accurate and consistent habitat, taxonomic and functional annotations of prokaryotic genomes. Nucleic Acids Res 2020;**48**(D1):D621-D625.
12. Wickham H. ggplot2: Elegant Graphics for Data Analysis. Springer-Verlag New York 2016.
13. Xiao L, Feng Q, Liang S, Sonne SB, Xia Z, Qiu X, Li X, Long H, Zhang J, Zhang D, Liu C, Fang Z, Chou J, Glanville J, Hao Q, Kotowska D, Colding C, Licht TR, Wu D, Yu J, Sung JJ, Liang Q, Li J, Jia H, Lan Z, Tremaroli V, Dworzynski P, Nielsen HB, Backhed F, Dore J, Le Chatelier E, Ehrlich SD, Lin JC, Arumugam M, Wang J, Madsen L, Kristiansen K. A catalog of the mouse gut metagenome. Nat Biotechnol 2015;**33**(10):1103-8.
14. Bergstrom A, Licht TR, Wilcks A, Andersen JB, Schmidt LR, Gronlund HA, Vigsnaes LK, Michaelsen KF, Bahl MI. Introducing GUt low-density array (GULDA): a validated approach for qPCR-based intestinal microbial community analysis. FEMS Microbiol Lett 2012;**337**(1):38-47.
15. Caesar R, Tremaroli V, Kovatcheva-Datchary P, Cani PD, Backhed F. Crosstalk between Gut Microbiota and Dietary Lipids Aggravates WAT Inflammation through TLR Signaling. Cell Metab 2015;**22**(4):658-68.
16. Jari Oksanen FGB, Michael Friendly, Roeland Kindt, Pierre Legendre, Dan McGlinn, Peter R. Minchin, R. B. O'Hara, Gavin L. Simpson, Peter Solymos, M. H. Stevens, Eduard Szoecs, Helene Wagner. vegan: Community Ecology Package. 2019;**R package version 2.5-6**.
17. Rogmann JJ. Ordinal Dominance Statistics (orddom): An R Project for Statistical Computing package to compute ordinal, nonparametric alternatives to mean comparison 2013;**Version 3.1.** .
18. Gregory R. Warnes BB, Lodewijk Bonebakker, Robert Gentleman, Wolfgang Huber, Andy Liaw, Thomas Lumley, Martin Maechler, Arni Magnusson, Steffen Moeller, Marc Schwartz and Bill Venables. gplots: Various R Programming Tools for Plotting Data. . 2020;**R package version 3.1.1.**
19. Hildebrand F, Tadeo R, Voigt AY, Bork P, Raes J. LotuS: an efficient and user-friendly OTU processing pipeline. Microbiome 2014;**2**(1):30.
20. McDonald D, Price MN, Goodrich J, Nawrocki EP, DeSantis TZ, Probst A, Andersen GL, Knight R, Hugenholtz P. An improved Greengenes taxonomy with explicit ranks for ecological and evolutionary analyses of bacteria and archaea. ISME J 2012;**6**(3):610-8.
21. Saary P, Forslund K, Bork P, Hildebrand F. RTK: efficient rarefaction analysis of large datasets. Bioinformatics 2017;**33**(16):2594-2595.
22. Kassambara A. ggpubr: 'ggplot2' Based Publication Ready Plots. 2020;**R package version 0.4.0**.
